# Multi-scale parameterization of neural rhythmicity with lagged Hilbert autocoherence

**DOI:** 10.1101/2024.12.05.627017

**Authors:** Siqi Zhang, Maciej J Szul, Sotirios Papadopoulos, Alice Massera, Holly Rayson, James J Bonaiuto

## Abstract

Analysis of neural activity in different frequency bands is ubiquitous in systems and cognitive neuroscience. Recent analytical breakthroughs and theoretical developments rely on phase maintenance of oscillatory signals without considering whether or not this assumption is met. Lagged (auto)coherence, the coherence between a signal and itself at increasing temporal delays, has been proposed as a way to quantify the rhythmicity, or periodicity, of a signal. However, current Fourier-based lagged autocoherence algorithms suffer from poor spectral accuracy and resolution, aliasing effects that become more pronounced at higher frequencies, and conflation with amplitude covariation, especially in frequency ranges in which the signal-to-noise ratio is low. We introduce a continuous estimator, lagged Hilbert autocoherence (LHaC), which addresses these limitations by using multiplication in the frequency domain for precise bandpass filtering, computing instantaneous analytic signals via the Hilbert transform, and thresholding using the amplitude covariation of phase-shuffled surrogate data. While LHaC and lagged Fourier autocoherence (LFaC) estimate distinct theoretical quantities, we compare their empirical behavior in simulations with controlled rhythmic structure. These analyses show that LHaC provides more spectrally resolved estimates of rhythmicity and is more sensitive to the duration of transient, short-lived oscillatory events. We further demonstrate the utility of LHaC for identifying frequency-specific differences in rhythmicity between conditions and tracking learning-related changes in neural oscillations. Lagged Hilbert autocoherence thus offers a refined and practically useful approach to characterizing neurophysiological rhythmicity.

## 1. Introduction

Neural oscillations are ubiquitous in the study of cognitive and systems neuroscience. The rhythmic activity of neuronal populations, spanning various frequency bands, forms a fundamental mechanism in the orchestration of large-scale brain network dynamics, influencing a multitude of cognitive and behavioral functions (Buzsáki and Freeman, 2015). Consequently, the accurate and reliable analysis of these oscillatory signals is paramount for advancing our understanding of brain function. The majority of theoretical developments to date have assumed the phase maintenance of oscillatory signals (Bonnefond et al., 2017; Fries, 2015; Jensen et al., 2015), however the veracity of this assumption is often overlooked in practice (Donoghue et al., 2022).

Traditional analyses have often inferred oscillatory properties from average signal energy (power spectra) or phase consistency across trials (e.g., inter-trial coherence). However, such measures do not directly quantify the persistence or periodicity of oscillations within single trials. To address this, lagged autocoherence has been proposed as a method to assess rhythmicity, defined as the degree to which a signal resembles a delayed copy of itself (Fransen et al., 2016, 2015). This approach captures the temporal autocorrelation of phase and amplitude structure over increasing lags, providing a direct index of rhythmic continuity. The most widely used implementation, lagged Fourier autocoherence (LFaC), has revealed meaningful patterns in alpha and beta rhythms across frequency bands, tasks, and populations (Fransen et al., 2016; Little et al., 2019; Rayson et al., 2023, 2022). However, LFaC is limited by poor spectral resolution, aliasing at higher frequencies, and confounding of rhythmicity with amplitude covariation under conditions with low signal-to-noise ratio (SNR) conditions. These issues arise from its reliance on short-time Fourier transforms and fixed-length windows, which impose tradeoffs between temporal and spectral resolution.

Recent theoretical and empirical work has also challenged the assumption that oscillatory signals maintain consistent phase or frequency over time (Donoghue et al., 2022). Transient, nonstationary bursts of rhythmic activity are increasingly recognized as an important mode of oscillatory dynamics (Sherman et al., 2016; van Ede et al., 2018; Rayson et al., 2022, 2023; Szul et al., 2023), necessitating tools that can track their time-varying structure with high frequency specificity. Improved spectral resolution is particularly important for identifying transient phenomena such as task-related beta and gamma bursts, or developmental shifts in alpha, effects that are easily obscured by imprecise spectral estimates.

In this context, we introduce a novel estimator, lagged Hilbert autocoherence (LHaC), which overcomes the limitations of LFaC by using bandpass filtering via frequency-domain multiplication, computing instantaneous analytic signals with the Hilbert transform, and thresholding against phase-shuffled surrogate data to control for amplitude covariation. While both LFaC and LHaC quantify lagged rhythmic structure, they estimate distinct theoretical quantities: LFaC relies on short-time Fourier-derived spectral power, whereas LHaC measures the lagged coherence of narrowband, continuous-time analytic signals. We compare their empirical behavior in simulations and in real EEG and MEG datasets. These analyses demonstrate that LHaC yields more spectrally resolved and behaviorally sensitive estimates of rhythmicity. In doing so, it enables new applications, including the detection of developmental trajectories in oscillatory dynamics and frequency-specific learning-related changes.

## 2. Methods

### 2.1 Lagged autocoherence algorithm

The original lagged autocoherence algorithm (Fransen et al., 2015) is based on a coherence metric commonly used to evaluate functional connectivity. Whereas coherence is typically applied to two different signals, lagged autocoherence operates on a single signal and a lagged copy of itself. The algorithm breaks the signal into evaluation windows, applies a Fourier transform to each window, and subsequently calculates coherence between temporally separated windows at a given frequency. The temporal gap between each window is specified in terms of cycles corresponding to the target frequency. The duration of each window can be set to match a prescribed number of cycles (e.g. 3 cycles no matter what the evaluated lag; Fransen et al., 2016, 2015; Little et al., 2019; Muralidharan et al., 2023), or to the evaluated lag (Rayson et al., 2022; Szul et al., 2023), and the windows can be either overlapping (Little et al., 2019; Rayson et al., 2022; Szul et al., 2023), or non-overlapping (Fransen et al., 2016, 2015; Muralidharan et al., 2023). Lagged autocoherence (*λ*) for a frequency *f* and lag *l*, is defined by:

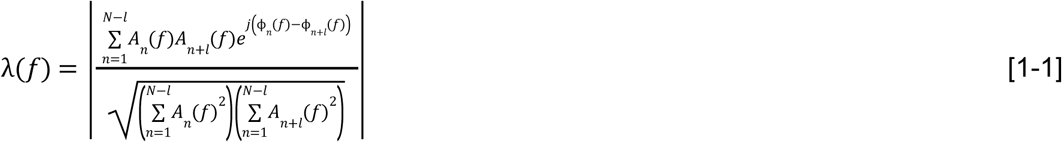

where 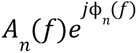 is the expression of the complex-valued Fourier coefficient of evaluation window *n* for frequency *f*, with amplitude *A* and phase *φ*. The Fourier coefficients are obtained by applying a Fourier transform to a Hann-tapered signal. The numerator, when divided by the number of window pairs, is called the lagged autospectrum. The lagged autospectrum depends on the difference in phase between windows pairs, but also on their amplitudes, and therefore it is normalized by the product of the sum of squared amplitudes from the first and second evaluation windows of each pair, which we refer to as the joint amplitude normalization factor. This results in a metric bounded by 0 and 1. Lagged Fourier autocoherence (LFaC) thus serves as a measure of phase synchrony between lagged segments of the signal at a specific frequency.

### 2.2 Lagged Hilbert autocoherence algorithm

Lagged Hilbert autocoherence (LHaC) diverges from the previously described lagged autocoherence algorithm by using Hilbert coherence (Bruña et al., 2018; Hu and Liang, 2012), and is therefore an adaptation of lagged Hilbert coherence (Pascual-Marqui et al., 2024). LHaC generates a continuous analytic signal using the Hilbert transform and then calculates coherence at a series of single time points corresponding to a specific lag duration. To compute LHaC at a given frequency and lag (in cycles of that frequency), the signal is first zero-padded and then bandpass-filtered by multiplication with a Gaussian kernel in the frequency domain (Figure 1a; see also Figure S1 for the kernel properties). The Gaussian is centered on the target frequency, and the kernel width (which is representative of the spectral width of the filter) can be appropriately adjusted. For single frequency analysis, we utilize a kernel width of 1 Hz. However, for a set of evenly spaced frequencies, the width is determined by the frequency resolution of the set. The resulting product is a bandpass filtered version of the signal. The Hilbert transform is subsequently applied to the bandpass filtered signal to yield the instantaneous amplitude, *A_*t*_*(*f*), and phase, ϕ*_*t*_*(*f*), of the signal at the specified frequency, *f*, for each time point, *t*. Following this, for a given signal recorded at a sampling rate *Fs*, lagged autocoherence is computed at time point pairs, starting from time point *s*, and separated in time by a delay, *d* (by time points), per the specified lag, *l*, and frequency 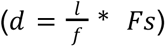:

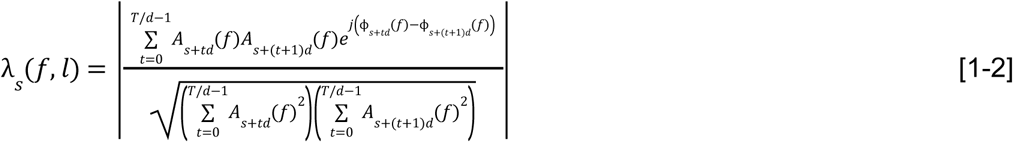

**Figure 1.**
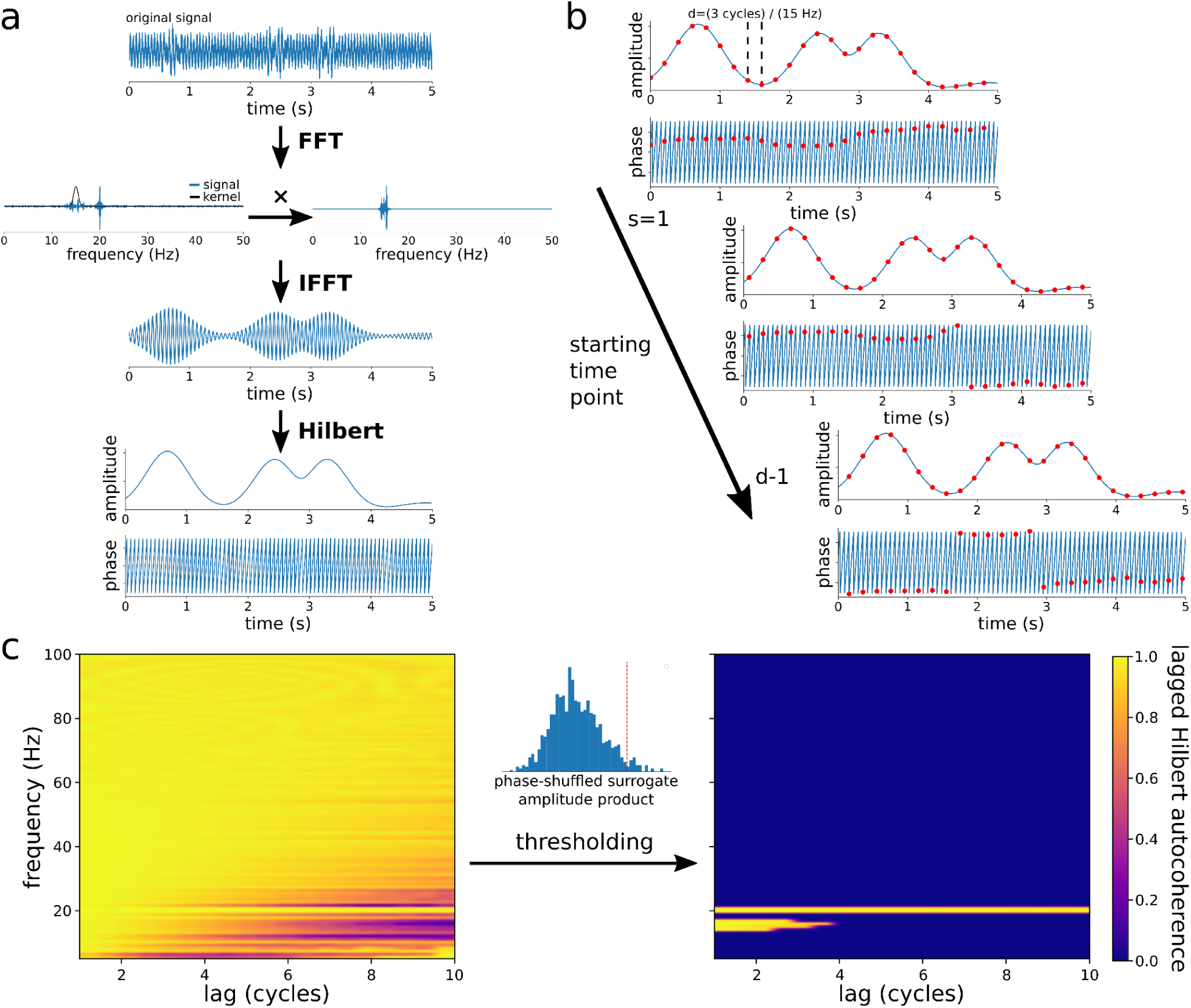
The lagged Hilbert autocoherence algorithm. a) The original time series is bandpass filtered at a given frequency using multiplication by a narrow Gaussian kernel in the frequency domain (centered around 15 Hz in this example) in order to obtain precise filtering, and then the Hilbert transform is used to obtain the instantaneous amplitude envelope and phase. b) Lagged autocoherence is computed using the instantaneous amplitude and phase between time steps separated by a given lag (evaluated at 15 Hz and 3 lag cycles in this example: d = (3 cycles / 15 Hz) ✕ Fs = 200 time points), and averaged over incremental shifts of the starting time point (as illustrated in equations [1-2] [1-3]). c) Lagged autocoherence is evaluated at a range of lags and frequencies. A threshold is computed from the average broadband amplitude product between time points of phase-shuffled surrogate data. Lagged Hilbert autocoherence is set to zero wherever the denominator is less than this threshold.

where *T* is the total number of time points. The starting time point, *s*, is initialized to the first time point in the series and is then shifted by one until all time points are incorporated (until *s* = *d* - 1), and the resulting lagged autocoherence values are averaged over all shifts (Figure 1b):

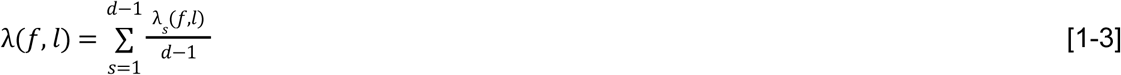

This approach yields a spectrally precise measure of lagged autocoherence; however, care must be taken when evaluating frequencies with low SNR. Narrow bandpass filtering in such regions can introduce spurious amplitude correlations, resulting in a very small denominator, the joint amplitude normalization factor, in equation [1-2]. When both the numerator and denominator approach zero, lagged autocoherence values can be artificially inflated, sometimes approaching 1 in the absence of genuine rhythmicity. This issue has been noted in prior work on coherence metrics (Cohen and Tsuchiya, 2018; Srinath and Ray, 2014), and highlights the need for a principled thresholding approach.

To avoid these artefacts, we use phase-shuffled surrogate data to estimate the distribution of spurious amplitude products under the null hypothesis of no phase structure. Specifically, for each trial, we apply phase randomization to the broadband signal: the Fourier transform is computed, the phase spectrum is randomized uniformly over [0, 2π] while preserving the amplitude spectrum, and the inverse transform yields a surrogate signal with the same broadband spectral content but no temporal phase structure. This surrogate is then used to compute the analytic signal and the amplitude product between adjacent time points. We repeat this procedure 1,000 times per trial. The 95th percentile of the resulting distribution of surrogate amplitude products for each trial is used as a threshold on the denominator of equation [1-2]. Any lagged autocoherence values computed from empirical data with a denominator below this threshold are set to zero (Figure 1c), reducing the likelihood of false positives in low-SNR conditions (see Figure S2 for threshold behavior across SNR levels). To validate this approach, we ran a simulation in which 15 Hz bursts were embedded in pink noise, and LHaC was evaluated at both 15 Hz and 50 Hz, a frequency with no underlying rhythmic signal. At 50 Hz, this mismatch produced artificially low values for the joint amplitude normalization factor and spurious inflation of LHaC values. As shown in Figure 2, the thresholding procedure effectively eliminated these false positives at lags greater than 1 cycle, confirming the necessity of joint amplitude normalization thresholding in preventing artefactual coherence under spectral mismatch.

**Figure 2.**
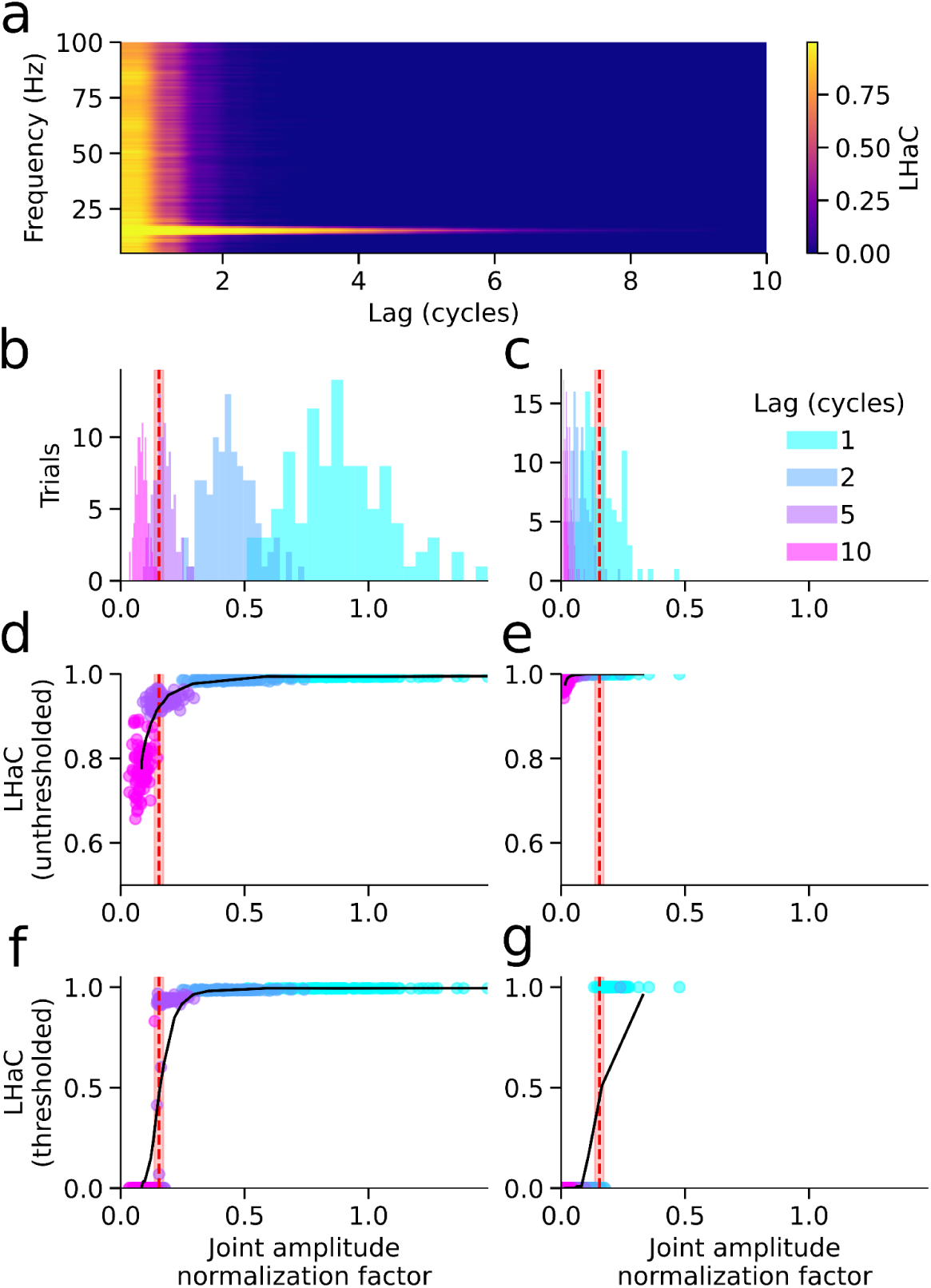
Joint amplitude normalization thresholding suppresses false-positive LHaC estimates. a) Mean LHaC as a function of lag (x-axis) and frequency (y-axis) for signals containing 15 Hz bursts embedded in pink noise (SNR = -10 dB). b-c) Distributions of the joint amplitude normalization factor (LHaC denominator) for signals evaluated at the true frequency (15 Hz, b) and at a mismatched frequency (50 Hz, c). Red dashed lines indicate the mean surrogate-based threshold; shaded regions indicate ±1 SD. Lag values of 1, 2, 5, and 10 cycles are shown in different colors. d-e) Unthresholded LHaC versus denominator for the true (d) and mismatched frequencies (e). f-g): Thresholded LHaC versus denominator for the true (f) and mismatched (g) frequency. Thresholding suppresses spuriously high LHaC values arising from low denominator values under spectral mismatch.

To assess whether the surrogate-based thresholding procedure controls false positives under the null hypothesis, we performed an additional simulation in which signals consisted solely of pink noise (1/f). Power spectra of both the original and phase-shuffled surrogate signals exhibited the expected 1/f profile (Figure 3a). We then evaluated LHaC at multiple frequencies and lags, applying our full pipeline including thresholding at α = 0.05. As shown in Figure 3b, the false positive rate at each frequency-lag combination remained below the nominal alpha level for lags ≥ 1 cycle, and declined to zero with increasing lag cycles. These results demonstrate that the surrogate-based thresholding procedure provides effective control of false positives under realistic colored noise conditions.

**Figure 3.**
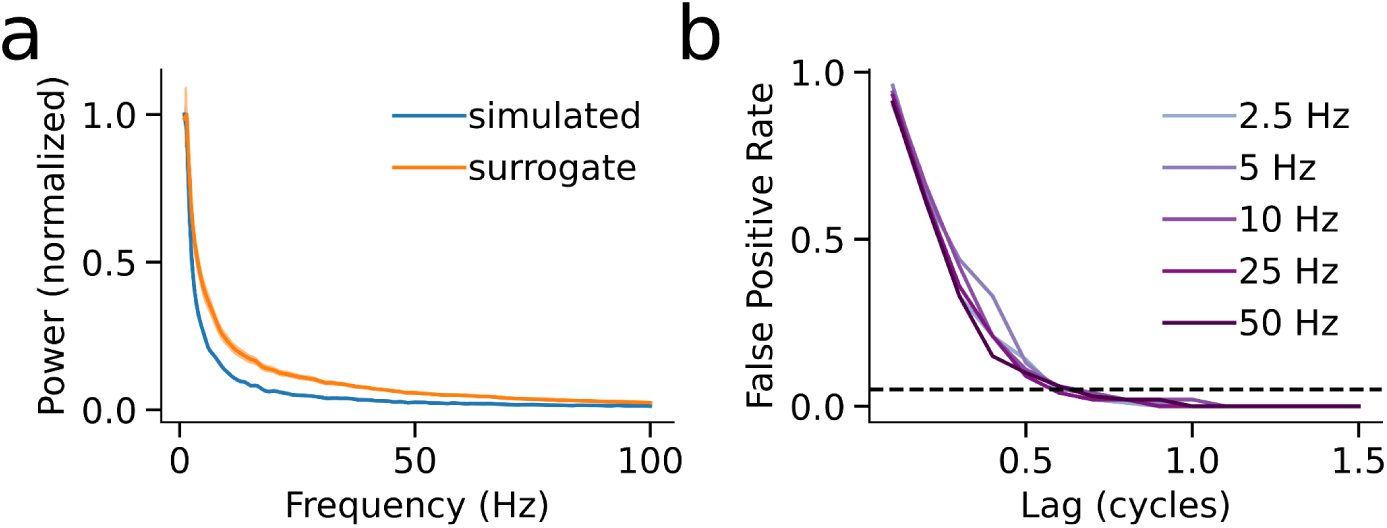
Surrogate-based thresholding controls false positives under the null hypothesis. a) Power spectra of simulated null data (pink noise) and corresponding phase-shuffled surrogates show matched 1/f profiles, indicating preserved spectral structure under phase randomization. Shaded regions denote ±1 SD across trials. b) False positive rate (FPR) as a function of lag, computed at five test frequencies. FPR was defined as the proportion of trials in which the joint amplitude normalization factor exceeded the 95th percentile of the surrogate distribution. For all test frequencies, FPR remained below the nominal alpha level (dashed line) for lags ≥ 1 cycle, demonstrating that the thresholding procedure effectively suppresses spurious detections under null conditions.

While both LFaC and LHaC can be used to characterize the rhythmic structure of finite neural signals, it is important to note that they are estimators of distinct population quantities. The underlying population definitions differ in how they incorporate lag and spectral information, and thus cannot be directly compared in terms of accuracy relative to a shared ground truth. Our comparisons below should therefore be interpreted as empirical analyses of their respective behavior in finite, noisy signals rather than evaluations of estimator performance in the statistical sense.

### 2.3 Simulations

To compare the spectral behavior of LFaC and LHaC under controlled conditions, we generated sinusoidal signals ranging from 10 to 50 Hz in increments of 5 Hz. For each frequency, 100 simulated trials were generated with a random initial phase for each trial. Trials had a duration of 5 s and a sampling rate of 500 Hz. The amplitude of the sinusoidal signals varied from -1 to 1 (au). To incorporate a realistic noise component, pink noise with a slope of 1 was introduced, scaled to achieve an SNR of -5 dB. PSDs for each simulated trial were computed using Welch’s method (Welch, 1967), with a 1 s window and 50% overlap. Lagged Fourier and Hilbert autocoherence were then computed across a range of 5 to 100 Hz, in 0.5 Hz increments, and from 1 to 6 cycles, in 0.5-cycle increments (see Figure S3 for an analysis of how LHaC results vary with lag sampling parameters). The normalized RMSE (as a fraction of total signal power) was computed between normalized PSDs and lagged autocoherence (averaged over lag cycles). Frequency spread was computed as the standard deviation of lagged autocoherence (averaged over lag cycles) over frequencies. For each metric, we fit a linear model with frequency, algorithm (LFaC versus LHaC), and their interaction as fixed effects. Fixed effects were assessed using type III F tests (car v3.1.0; Fox et al., 2019), and post-hoc comparisons between algorithms were performed using estimated marginal means (emmeans v1.10.0; Lenth, 2024). Differences in slope across frequency were tested using pairwise comparisons of simple trends. We repeated these simulations at five SNRs levels (-50, -20, -15, -5, 0 dB) and analyzed normalized RMSE and frequency spread at each level.

To determine the sensitivity of LHaC and LFaC to burst duration and the number of bursts, we ran two sets of simulations, each with 400 simulated trials of 10 s with a 500 Hz sampling rate and an SNR of -5 dB. Both sets of simulations involved generating signals with bursts of oscillatory activity at frequencies ranging from 20 to 50 Hz, in increments of 5 Hz. Because our simulated signals consisted of transient bursts of oscillatory activity, the amplitude envelope exhibited strong short-term autocovariance. These dynamics are known to impair the interpretability of conventional coherence metrics, providing a relevant testbed for evaluating estimator behavior under nonstationary amplitude structure. The first set of simulations generated one burst per trial, with a random duration from 2 to 5 cycles in each trial. In the second set of simulations, there were 1 to 5 bursts in each trial, each 3 cycles long. In both sets of simulations, LHaC and LFaC were evaluated from 5 to 100 Hz in 0.5 Hz increments, and from 0.25 to 20 cycles, in increments of 0.25 cycles. The decrease in autocoherence with increasing lags was then evaluated by fitting an inverse sigmoid function to the LHaC and LFaC at the simulated burst frequency over all evaluated lag cycles:

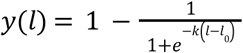

where *l* is the evaluated lag cycle, and the fitted variables *k* and *l_0_* determine the rate of autocoherence decay with increasing lags, and the crossover point in terms of lag cycles corresponding to the greatest lagged autocoherence decrease, respectively. Linear models were used to evaluate the relationship between the fitted values of *k* and *l_0_*and the simulated burst duration or number. Each of these models had *k* or *l_0_* as the dependent variable, and frequency, a single burst metric, and their interaction as fixed effects. The burst metrics used were burst duration in seconds, burst duration in cycles, and the number of bursts. Fixed effects were assessed using type III F tests (car v3.1.0; Fox et al., 2019), and for each inverse sigmoid parameter, models with burst duration in seconds or cycles were compared using the Akaike information criterion (AIC; Akaike, 1974, 1973). We further repeated all burst simulations at five SNRs (-50, -20, -15, -5, 0 dB) and examined how the slope of burst duration or number effects on *k* and *l_0_*varied with SNR.

While we compare the empirical behavior of the LFaC and LHaC estimators, we do not define a shared population lagged autocoherence function, nor do we simulate from a stochastic model with a known parametric form. Therefore, these simulations do not constitute formal comparisons of estimator performance in the classical statistical sense. Instead, we evaluate how well each method recovers key structural features of the signal, such as frequency content and lag-specific decay, when applied to signals with known oscillatory properties embedded in colored noise.

### 2.4 Datasets and Applications

After establishing the utility of lagged Hilbert autocoherence and contrasting it with lagged Fourier autocoherence in simulation, we applied both algorithms to two different representative empirical datasets: EEG recorded during action observation and execution in infants and adults, and MEG recorded during visuomotor adaptation in adults. These modalities, widely used in human cognitive neuroscience, offer complementary strengths in temporal and spatial resolution and serve to illustrate the practical value of LHaC across diverse experimental contexts.

#### 2.4.1 EEG data

EEG data were previously recorded from human infants and adults as part of a study investigating the neural bases of action execution and observation (methodological details can be found in Cannon et al., 2014; Yoo et al., 2016). The experiment was approved by the University of Maryland Institutional Review Board, with informed consent obtained from the infants’ parents and adult participants. After exclusions, the sample included 28 9-month-old infants, 33 12-month-old infants, and 22 adults. Infant and adult participants performed the same reach-and-grasp task. Infants were seated on their caregiver’s lap and adults in a chair during an observation and execution condition. In the observation trials, participants watched a female presenter interact with a toy using a hand-operated tool, while in the execution trials, participants were given the opportunity to reach for the toy themselves. EEG data were recorded using a 65-channel HydroCel Geodesic Sensor Net (Electrical Geodesics, Inc., Eugene, OR), sampled at 500 Hz, with the vertex (Cz) electrode used as an online reference, and impedances kept below 100 kΩ. The EEG data were later exported to a MATLAB compatible format for offline processing.

Data preprocessing was conducted in MATLAB R2018a, using a modified MADE pipeline (Debnath et al., 2020) with artifact detection routines from the NEAR pipeline (Kumaravel et al., 2022). Signals were high-pass filtered at 1 Hz and low-pass filtered at 100 Hz. Artifact-laden channels were identified and removed with the NEAR pipeline’s local outlier factor metric, followed by artifact subspace reconstruction for non-stereotyped artifacts. Independent component analysis (ICA) was performed on 1s epochs to detect and remove stereotyped artifacts like eye blinks, with artifactual components identified via the Adjusted-ADJUST plugin (Leach et al., 2020; Mognon et al., 2011). Behavioral events were captured via video recording, and for both conditions, five coded time points were used to segment the EEG time series into 3.5s epochs centered on the baseline period, the go cue, the first touch with the hand, the completion of the grasp, and the end of the movement. Epochs exceeding 10% artefacted channels were excluded; otherwise, artefacted channels were interpolated. Missing channels were subsequently interpolated, data were average re-referenced, and line noise (60 Hz) was removed using the Zapline algorithm (de Cheveigné, 2020). Subjects with fewer than five trials for any condition epoch were excluded, resulting in a final sample of 21 9-month-olds, 24 12-month-olds, and 22 adults.

Lagged Hilbert and Fourier autocoherence was computed for each participant in both the observation and execution conditions, and in each epoch, from 4 to 100Hz in 0.5 Hz increments and 0.1-4.5 cycles in increments of 0.1 cycles. The lagged autocoherence values were averaged over all electrodes in C3 and C4 clusters (E16, E20, E21, E22, E41, E49, E50, E51). To isolate the periodic spectral component for comparison with lagged autocoherence, we first computed power spectral densities (PSDs) using Welch’s method with 1 s Hann windows and 50% overlap. We then applied the specparam algorithm (Donoghue et al., 2020) to each PSD in order to decompose the spectrum into aperiodic and periodic components (peak width limits = 1-8 Hz and a maximum of 6 peaks). The periodic component was derived by subtracting the aperiodic component from the PSD. To explore the interacting effects of condition and epoch on LHaC and LFaC, we applied a mass-univariate two-way repeated measures ANOVA (2 conditions by 5 epoch levels; including the baseline epoch) within each age group, and follow-up mass-univariate t-tests to explore significant interactions between condition and epoch. All of these analyses were corrected for multiple comparisons at the cluster level.

To assess whether condition-related differences in alpha-band LHaC reflected changes in oscillatory burst structure, we quantified alpha burst duration separately for each age group using age-specific alpha ranges determined from the group-level periodic spectra (9 months: 3.5 - 9 Hz; 12 months: 6 - 9 Hz; adults: 9 - 13 Hz). EEG data were band-pass filtered within the relevant alpha range and the analytic signal was computed via the Hilbert transform to obtain the instantaneous amplitude envelope. Bursts were defined as contiguous periods where the amplitude exceeded a dynamic threshold set to the 75th percentile of the envelope across the target trial and its eight surrounding trials. Burst duration was computed both in cycles and in milliseconds. Analyses were restricted to electrodes over the same C3 and C4 electrode clusters used in the LHaC and LFaC analyses. For each age group, linear mixed models were used to analyze burst duration, in cycles and in milliseconds (absolute duration), with condition (execution versus observation), epoch (baseline, go, first touch, grasp completion, movement end), and their interaction as fixed effects, and by-subject random intercepts and slopes for condition and epoch (lmer v1.1-31; Bates et al., 2015). Fixed effects were evaluated using type III Wald chi-square tests (car v3.1.0; Fox & Weisberg, 2019), and pairwise comparisons were computed using estimated marginal means (emmeans v1.10.1; Lenth, 2024).

#### 2.4.2 MEG data

High precision MEG data were recorded for a previous experiment from 38 human adults (25 female) as they performed a visually cued button press task (methodological details can be found in the corresponding manuscript; Szul et al., 2023). The study protocol was in accordance with the Declaration of Helsinki, and all participants gave written informed consent which was approved by the regional ethics committee for human research (CPP Est IV - 2019-A01604-53). The data were recorded using a 275-channel Canadian Thin Films (CTF) MEG system featuring SQUID-based axial gradiometers (CTF MEG Neuro Innovations, Inc. Coquitlam, Canada) inside a magnetically shielded room. Visual stimuli were displayed via a projector on a screen roughly 80 cm from the participant, and a joystick (NATA Technologies, Canada) was used for participant responses. The MEG data collected were digitized continuously at a sampling rate of 1200 Hz. Participants performed a cued visuomotor adaptation task, making rapid joystick-based movements to reach visually presented targets, guided by a random dot kinematogram (RDK). In some trials, the position of the cursor indicating the joystick position was rotated by 30°. Participants were divided into two groups, explicit (*N* = 20) and implicit (*N* = 18). The direction of the explicit group’s visuomotor rotation was predicted by the RDK’s direction of coherent motion, while the reaches of the implicit group were subject to a constant rotation unrelated to the RDK. Participants underwent training blocks, followed by one block of trials without visuomotor rotation, seven blocks of trials with visuomotor rotation, and a final washout block without rotation. The MEG data was preprocessed using the MNE-Python toolbox (Gramfort et al., 2014), downsampled to 600 Hz, and filtered with a low pass 120 Hz zero-phase FIR filter. Line noise was removed using an iterative version of the Zapline algorithm (de Cheveigné, 2020). Ocular and cardiac artifacts were identified and removed using Independent Component Analysis, correlating components with eye movement signals and ECG R peak detection, respectively.

The data was then epoched around the visual stimulus onset (-1 to 2 s) and the end of the reaching movement (-1 to 1.5 s), focusing on 11 sensors above the left sensorimotor area. One subject was excluded from further analysis due to a technical error during recording, and five subjects were excluded because their power spectra did not contain any periodic peaks (final implicit *N* = 18, explicit *N* = 14). Lagged Hilbert autocoherence and lagged Fourier autocoherence were computed for each epoch of each trial from 4 to 100 Hz in increments of 0.5 Hz and from 0.1 to 4.5 lag cycles in increments of 0.1 cycles. For each subject, lagged autocoherence values were then averaged over channels and then over trials within each block and normalized according to the maximal lagged autocoherence in the first block. PSDs were computed using Welch’s method with 1 s Hann windows and 50% overlap, and the specparam algorithm (Donoghue et al., 2020) was used to decompose each PSD into aperiodic and periodic components (peak width limits = 1-8 Hz and a maximum of 6 peaks). Within the alpha (7-13 Hz) and beta (15-30 Hz) bands, we fitted an inverse sigmoid to the LHaC and LFaC curves across lag cycles and extracted the crossover point and decay rate for each block. These parameters were analyzed relative to the baseline (first) block. For each block, these values were compared between groups using a bootstrap permutation test. This involved generating null distributions (N = 10,000) by sampling participants with replacement and randomly assigning them to a group before computing the group difference.

To complement the permutation-based group comparisons, we applied linear mixed-effects models (lmer v1.1-31; Bates et al., 2015) to examine how crossover point and decay rate varied with group and block structure across learning. For each algorithm, frequency band, and task epoch, we first fit a model with group, block type (baseline, rotation, washout), and their interaction as fixed effects, and subject as a random intercept. Fixed effects were evaluated using type III Wald chi-square tests (car v3.1.0; Fox & Weisberg, 2019), and pairwise comparisons were computed using estimated marginal means (emmeans v1.10.1; Lenth, 2024). To test for linear trends in rhythmicity within the adaptation phase, we fit a second linear mixed model restricted to the rotation blocks, with group, block (as a continuous variable), and their interaction as fixed effects, again with subject as a random intercept. We followed up significant interactions using estimated marginal trends (emmeans v1.10.1; Lenth, 2024) to compare linear slopes between groups. This allowed us to assess not only condition differences and block-type effects, but also the extent to which rhythmicity parameters changed systematically over the course of adaptation.

## 3. Results

### 3.1 Simulation comparison of lagged Fourier autocoherence and lagged Hilbert autocoherence

We first sought to validate the performance of lagged Hilbert autocoherence (LHaC) by comparing it to lagged Fourier autocoherence (LFaC) in controlled simulations. These simulations were designed to assess three core properties of the estimators: (1) spectral accuracy, quantified by the normalized root mean square error (RMSE) between lagged autocoherence, averaged over lags, and the power spectral density (PSD); (2) spectral precision, indexed by the frequency spread (standard deviation over frequency); and (3) temporal sensitivity to rhythmic structure, assessed via parametric fits of inverse sigmoid functions to characterize the decay of coherence across lag cycles. We varied the simulated signal properties across frequency, burst duration, burst count, and SNR, and compared how LFaC and LHaC responded to each manipulation. These simulations enabled us to empirically characterize the resolution, robustness, and interpretability of both metrics across diverse spectral and temporal regimes.

To characterize the empirical spectral accuracy and precision of LHaC and LFaC, we simulated oscillatory neural signals at frequencies from 10 to 50 Hz, ran LHaC and LFaC on the simulated signals, and then averaged each lagged autocoherence metric over simulated trials and then over 1 to 6 lag cycles. We then computed the RMSE between normalized PSDs of the simulated signal and lagged autocoherence, and the frequency spread (standard deviation over frequencies) of lagged autocoherence. For interpretability, we express RMSE as a fraction of the total power of the signal, yielding a normalized RMSE. The RMSE quantifies the similarity between the lagged autocoherence and the PSD-derived spectral profile, serving as a practical index of empirical agreement. We use this as a heuristic to characterize differences in spectral resolution between LFaC and LHaC, while acknowledging that they estimate distinct theoretical quantities. The frequency spread indicates the precision of the lagged autocoherence measure over frequencies, with lower values reflecting a sharper and more precise spectral profile. Across simulations, we found that the Hilbert-based lagged autocoherence exhibited closer empirical correspondence to the simulated PSD than LFaC, with lower RMSE relative to the PSD of the simulated signal across all tested frequencies (Figure 4a-d; main effect of algorithm: *F*(1,1796) = 614.60, *p* < 0.001). The RMSE slope across frequency also differed significantly between algorithms, decreasing for LHaC and increasing for LFaC (slope difference: *t*(1796) = 58.22, *p* < 0.001). LHaC also had greater spectral precision, with significantly lower frequency spread than LFaC (Figure 4a-c,e; main effect of algorithm: *F*(1,1796) = 5151.56, *p* < 0.001), and a significantly shallower slope across frequency (*t*(1796) = 15.95, *p* < 0.001).

**Figure 4.**
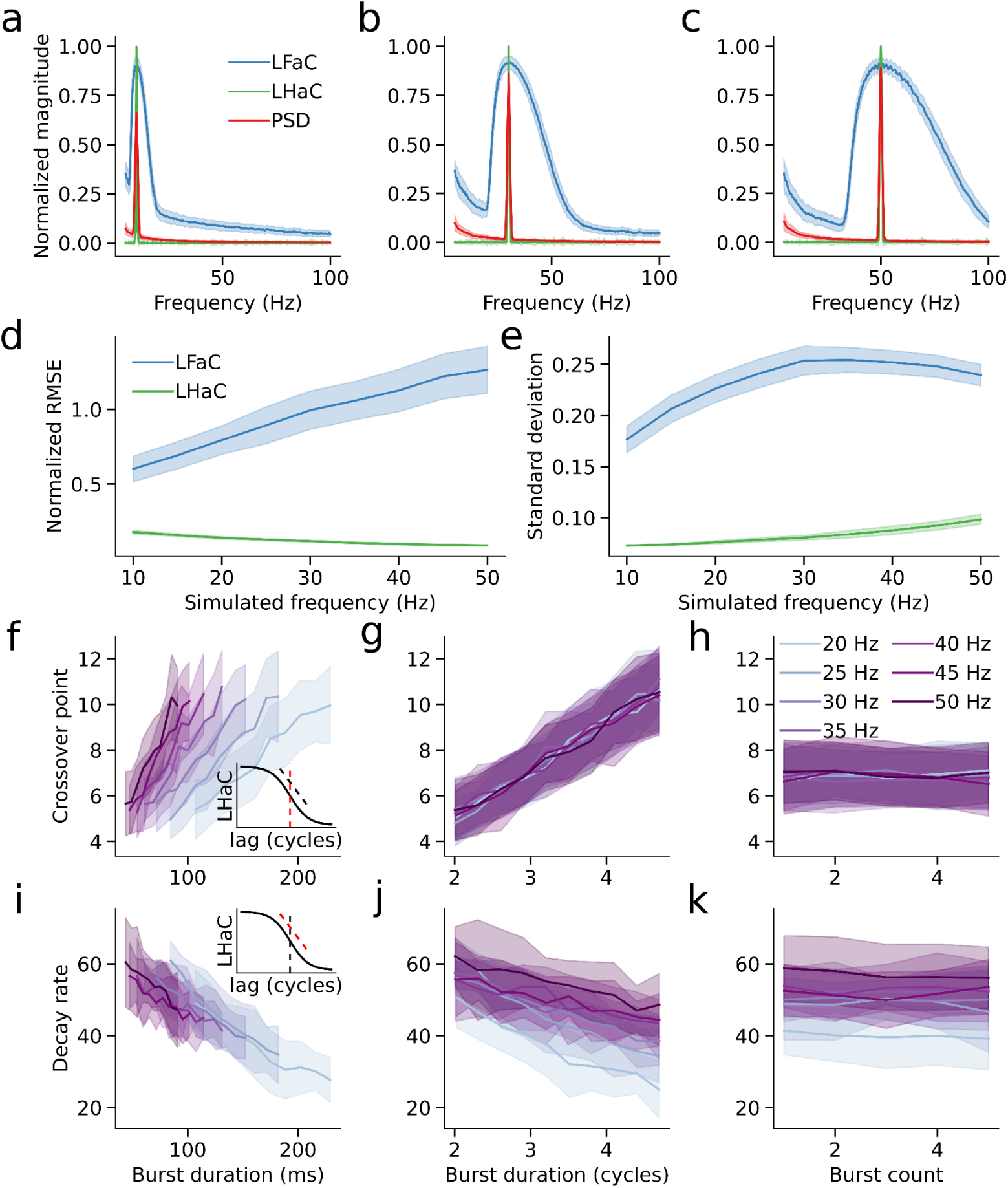
Lagged Fourier autocoherence and lagged Hilbert autocoherence simulation results. Lagged Fourier autocoherence and lagged Hilbert autocoherence averaged over 1 to 6 cycles and normalized, compared to the normalized PSD for simulations of oscillations at 10 (a), 25 (b), and 50 Hz (c). d) Normalized RMSE of the averaged and normalized LFaC and LHaC metrics compared to the normalized PSD over simulated oscillations from 10 to 50Hz. e) As in (d), for the standard deviation over frequencies of LFaC and LHaC, averaged over 1 to 6 cycles. f) For each simulated burst frequency, the crossover point of the inverse sigmoid fitted to LHaC (inset: red dashed line) as a function of burst duration in seconds. The solid lines denote the mean within 10 equally spaced bins, and the shaded area represents the standard deviation. g) As in (f), with burst duration expressed in terms of the number of cycles of the simulated burst frequency. h) As in (f), for the number of simulated bursts. i) For each simulated burst frequency, the decay rate of the inverse sigmoid fitted to LHaC (inset: red dashed line) as a function of burst duration in seconds. The solid lines denote the mean within 10 equally spaced bins, and the shaded area represents the standard deviation. j) As in (i), with burst duration expressed in cycles. k) As in (i), for the number of simulated bursts.

Having characterized the empirical spectral resolution and precision of LHaC relative to LFaC, we next examined how LHaC behaves as a function of burst duration and burst count over increasing lag cycles. We therefore simulated bursts of oscillatory activity, spanning frequencies of 20 to 50 Hz. In the first simulation set, each trial was characterized by a single burst whose duration was uniformly randomized between 2 and 5 cycles. Conversely, in the subsequent set, every burst lasted for 3 cycles, but each trial consisted of 1 to 5 bursts (with a uniform random distribution over trials). We fit an inverse sigmoid function to LHaC at the simulated frequency, yielding both a crossover point (Figure 4f, inset) and a decay rate (Figure 4i, inset), representing the number of lag cycles corresponding to the maximal decrease in lagged autocoherence and the rate of this decrease, respectively. Because the crossover point is expressed in lag cycles, it should scale with burst duration when measured in cycles, whereas the decay rate, having units of lagged autocoherence per cycle, should more closely reflect a temporal decay constant and therefore align with burst duration in milliseconds.

The fitted values for both the crossover point and decay rate were then compared against the burst duration and the number of bursts for every simulated frequency. This revealed a strong linear relationship between the inverse sigmoid crossover point and burst duration when measured in cycles (*F*(1) = 333.37, *p* < 0.001), with no significant interaction with frequency (*F*(1) = 0.22, *p* = 0.64; Figure 4g). While expressing burst duration in milliseconds yielded a significant interaction with frequency (*F*(1) = 733.98, *p* < 0.001; Figure 4f), model comparison showed no meaningful difference between models using milliseconds or cycles (ΔAIC = -0.18). In a contrasting pattern, the decay rate of the inverse sigmoid had an inverse relationship with burst duration (milliseconds: *F*(1) = 215.00, *p* < 0.001; cycles: *F*(1) = 408.23, *p* < 0.001). However, the offset of this relationship varied with frequency when burst duration was expressed in cycles (Figure 4j; *F*(1) = 109.81, *p* < 0.001), but was more stable across frequencies when duration was expressed in milliseconds (Figure 4i; *F*(1) = 6.12, *p* = 0.013). Model comparison favored the formulation based on seconds (*ΔAIC* = -2927.07), indicating that absolute duration provides a more parsimonious explanation of the decay rate pattern across frequencies. Notably, the number of bursts had no significant effect on the decay rate of the inverse sigmoid (Figure 4k; *F*(1) = 0.48, *p* = 0.487), nor did it interact with frequency (*F*(1) = 0.05, *p* = 0.827). For the crossover point, there was a marginal interaction with frequency (*F*(1) = 3.86, *p* = 0.050; Figure 4h), but it did not survive correction for multiple comparisons. LHaC therefore provides a robust estimate of burst duration that is insensitive to burst count.

To evaluate how signal-to-noise ratio (SNR) influences the empirical spectral properties of LHaC and LFaC, we repeated all simulations across five SNR levels: -50, -20, -15, -5, and 0 dB. We assessed spectral profile similarity using normalized RMSE, and spectral precision using frequency spread (standard deviation of the autospectrum). Across all SNR levels, LHaC consistently exhibited lower RMSE than LFaC (all *p* < 0.001; Figure 5a), reflecting closer empirical alignment with the simulated spectral content even under high noise. The RMSE gap widened with increasing SNR, reflected in a significant interaction between frequency and algorithm from -20 dB upward (e.g., -5 dB: *F*(1,1796) = 3389.84, *p* < 0.001). While RMSE slopes were similar at low SNR (-50 dB: *t*(1796) = -1.34, *p* = 0.18), a divergence emerged at higher SNRs, with LHaC trending downward and LFaC increasing with frequency (0 dB: slope difference *t*(1796) = 101.18, *p* < 0.001).

**Figure 5.**
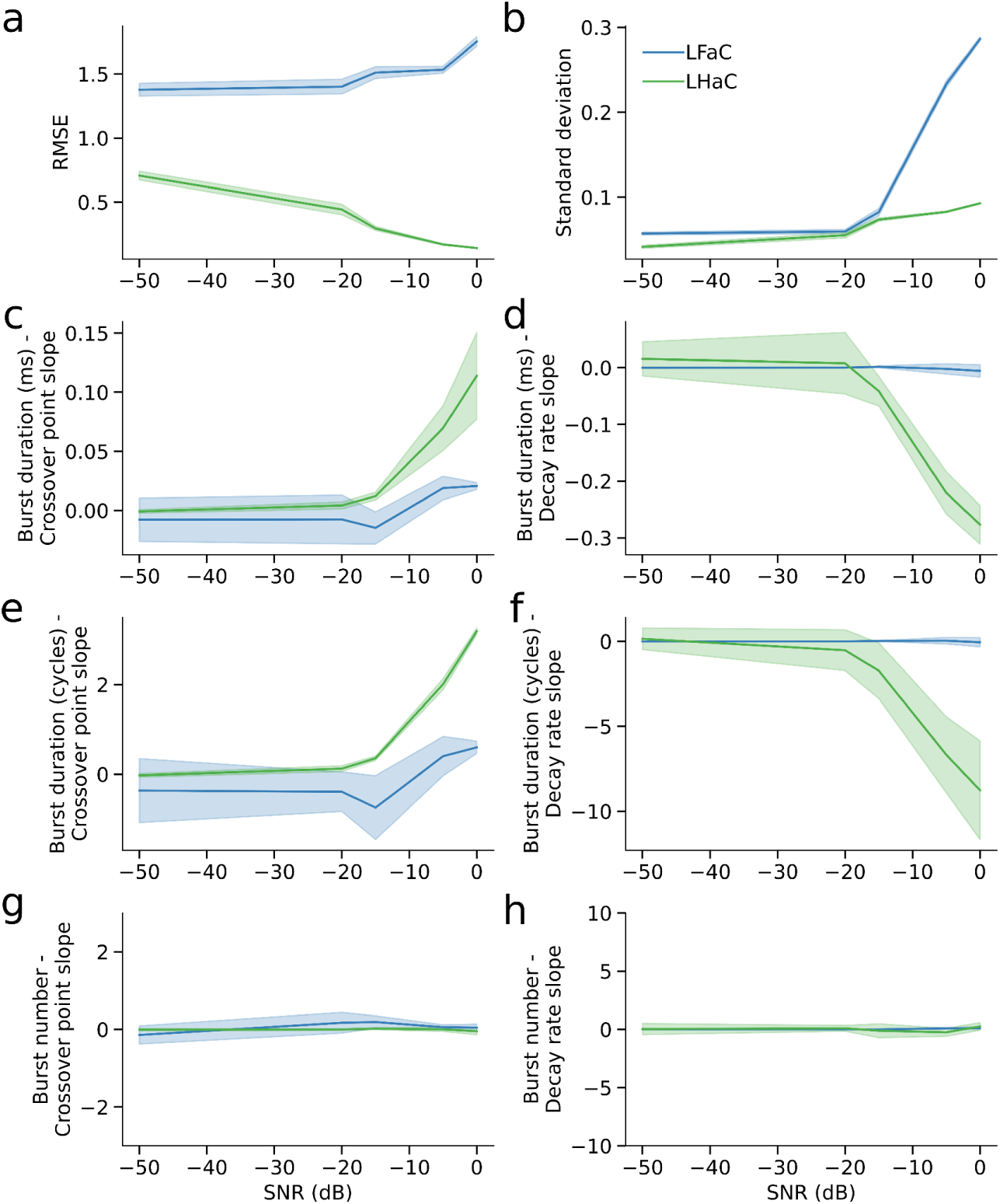
Lagged Hilbert autocoherence maintains spectral fidelity and sensitivity to burst dynamics across a range of signal-to-noise ratios. a-b) Compared to LFaC, LHaC consistently exhibited lower normalized root mean squared error (RMSE; a) and reduced frequency spread (standard deviation; b) relative to the power spectra of the simulated signals across SNR levels. c-d) At higher SNRs, the magnitude of the slope of the relationship between burst duration (in milliseconds) and the crossover point (c) and decay rate (d) increased for LHaC, but not LFaC. e-f) When burst duration was measured in cycles, the crossover point slope (e) and decay rate slope (f) for LHaC remained robust across SNRs, while LFaC again showed minimal sensitivity. g-h) Burst number had weak or negligible effects on crossover point (g) and decay rate (h) slopes at all SNR levels, with LHaC showing only marginal trends and LFaC showing none. Shaded areas denote ±1 SD across simulated frequencies.

LHaC also provided more spectrally precise estimates across all SNRs, as reflected by consistently lower frequency spread compared to LFaC (*p* < 0.001 at each SNR; Figure 5b). While the difference in spread was modest at -50 dB (*t*(1796) = 57.14, *p* < 0.001), it increased with SNR, reaching a five-fold difference at 0 dB (*t*(1796) = 246.15, *p* < 0.001). The slope of frequency spread across frequency also differed significantly between estimators above -20 dB, with LHaC showing more stable or shallower increases (0 dB: slope difference *t*(1796) = -3.64, *p* = 0.0003). Taken together, these results indicate that LHaC exhibited a closer empirical correspondence to the spectral content of the simulated signal than LFaC across a range of SNRs, and that this became more pronounced under moderate to high SNR conditions.

We next examined whether the relationship between burst duration and lagged autocoherence crossover point and decay rate was robust across SNR levels (Figures S4-5). At low SNR (-50 and -20 dB), burst duration had minimal or inconsistent effects on the LHaC crossover point and decay rate, with weak model fits and ΔAIC values near zero. However, as SNR increased (-15 to 0 dB), clearer relationships emerged: the LHaC crossover point became increasingly well predicted by burst duration in cycles rather than milliseconds (e.g., 0 dB: *F*(1) = 653.37, *p* < 0.001; ΔAIC = -2.65), whereas the decay rate was better explained by burst duration in milliseconds (e.g., 0 dB: *F*(1) = 838.75, *p* < 0.001; ΔAIC = -450.65; Figure 5c-f). These findings indicate that, at sufficient SNR, the LHaC crossover point reflects burst duration in cycles, whereas the decay rate reflects the absolute burst duration. By contrast, the number of bursts had inconsistent effects across SNR levels (Figure 5g,h; Figure S6). At -50 dB, burst number significantly influenced the LHaC crossover point (*F*(1) = 7.87, *p* = 0.005), but not the decay rate (*p* = 0.33). These effects diminished at higher SNRs: above -20 dB, neither parameter showed reliable associations with burst number (all *p* > 0.05). Thus, under realistic noise conditions, LHaC parameters remain primarily sensitive to burst duration rather than burst rate. These relationships were either extremely weak or entirely absent with LFaC. Across all SNR levels, neither burst duration nor number reliably predicted the crossover point or decay rate derived from LFaC (Figure 5c-h), indicating that the metric lacks sensitivity to the temporal structure of transient oscillatory events.

### 3.2 Sensorimotor rhythmicity is differentially modulated during action observation and execution conditions

We next sought to evaluate the practical utility of LHaC for detecting frequency-specific rhythmic differences across conditions and developmental stages using EEG data. Specifically, we tested whether LHaC could (1) capture developmental changes in rhythmicity during action execution and observation in infants and adults, and (2) resolve condition- and epoch-specific modulations in neural rhythmicity that differ between groups. To benchmark its performance, we compared LHaC with LFaC using three criteria: empirical alignment with the signal’s spectral content (normalized RMSE to the PSD), frequency-band specificity (localization in frequency-lag space), and sensitivity to experimental manipulations (condition × epoch interactions). This dataset allowed us to assess the resolution and sensitivity of lagged autocoherence metrics in a naturalistic sensorimotor context with rich spectral structure, where infant alpha rhythms are known to differ from those of adults. By contrasting observation and execution conditions across three age groups (9-month-olds, 12-month-olds, adults), we examined whether LHaC can reliably capture group- and condition-dependent dynamics of alpha, beta, and gamma rhythmicity across distinct sensorimotor epochs.

We applied LHaC and LFaC to EEG signals recorded over sensorimotor cortex in 9- and 12-month-old infants and adults during both execution and observation of grasping a toy (Cannon et al., 2014; Yoo et al., 2016). The within-group averaged LHaC across conditions and epochs of 9-month (Figure 6a), 12-month (Figure 6b), and adult group (Figure 6c) exhibited peak values within the theta, alpha, beta, and gamma frequency ranges at low lags (Figure 6d-f), with peak frequency increasing between infancy and adulthood. In the two infant groups, the alpha-band peak occurred around 8 Hz, but differed in decay rate, while in the adult group (Figure 6c), the alpha-band peak appeared around 12 Hz and exhibited a slower decay, suggesting developmental increases in both alpha peak frequency and rhythmicity. In contrast, only the alpha peaks were prominent in LFaC (Figure 7a-c). Comparison of normalized RMSE relative to the periodic component of the PSD revealed lower RMSE for LHaC than LFaC across all age groups (9 months: LFaC = 0.206, LHaC = 0.105; 12 months: LFaC = 0.298, LHaC = 0.093; adults: LFaC = 0.575, LHaC = 0.094), indicating closer empirical alignment between LHaC and spectral features of the signal across development.

**Figure 6.**
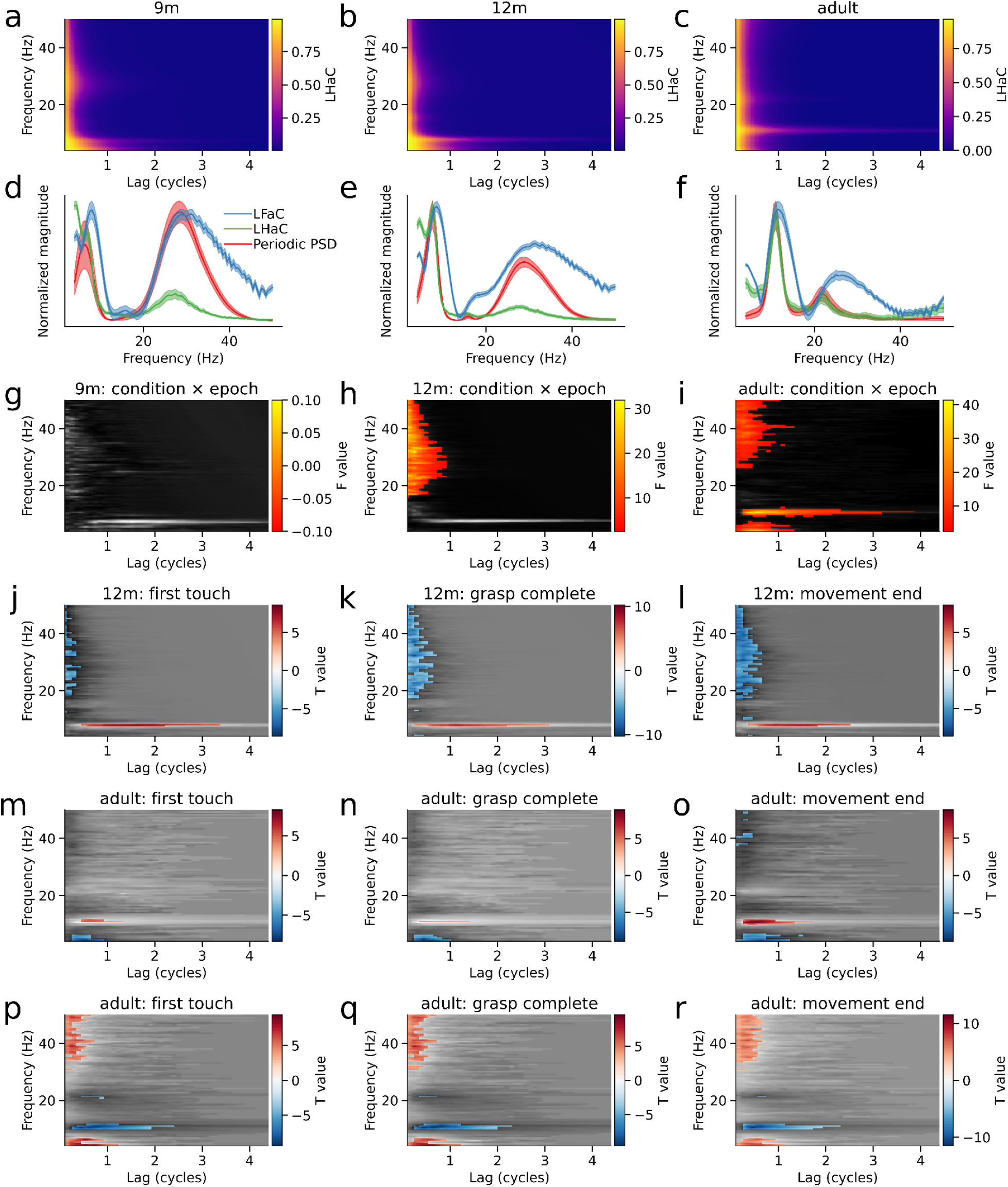
Sensorimotor rhythmicity differs between action execution and observation. a-c) Group-averaged lagged Hilbert autocoherence (LHaC) computed within EEG electrode clusters over sensorimotor cortex from 9-month infants (a), 12-month infants (b), and adults (c). Alpha peak frequency and rhythmicity increases with age. d-f) Normalized periodic PSD components, LHaC, and LFaC (averaged over lags) from the same electrode clusters for each group. g-i) LHaC F statistics for the condition by epoch interaction for each group. Statistically significant clusters, corrected using cluster-based permutation tests, are shown in color. j-l) LHaC T-statistics for epochs (aligned to the first touch, grasp completion, and end of the movement) with between-condition differences for the 12-month infants. m-o) and p-r) LHaC T-statistics for epochs with between-condition and between-epoch differences, respectively, for the adult group.

**Figure 7.**
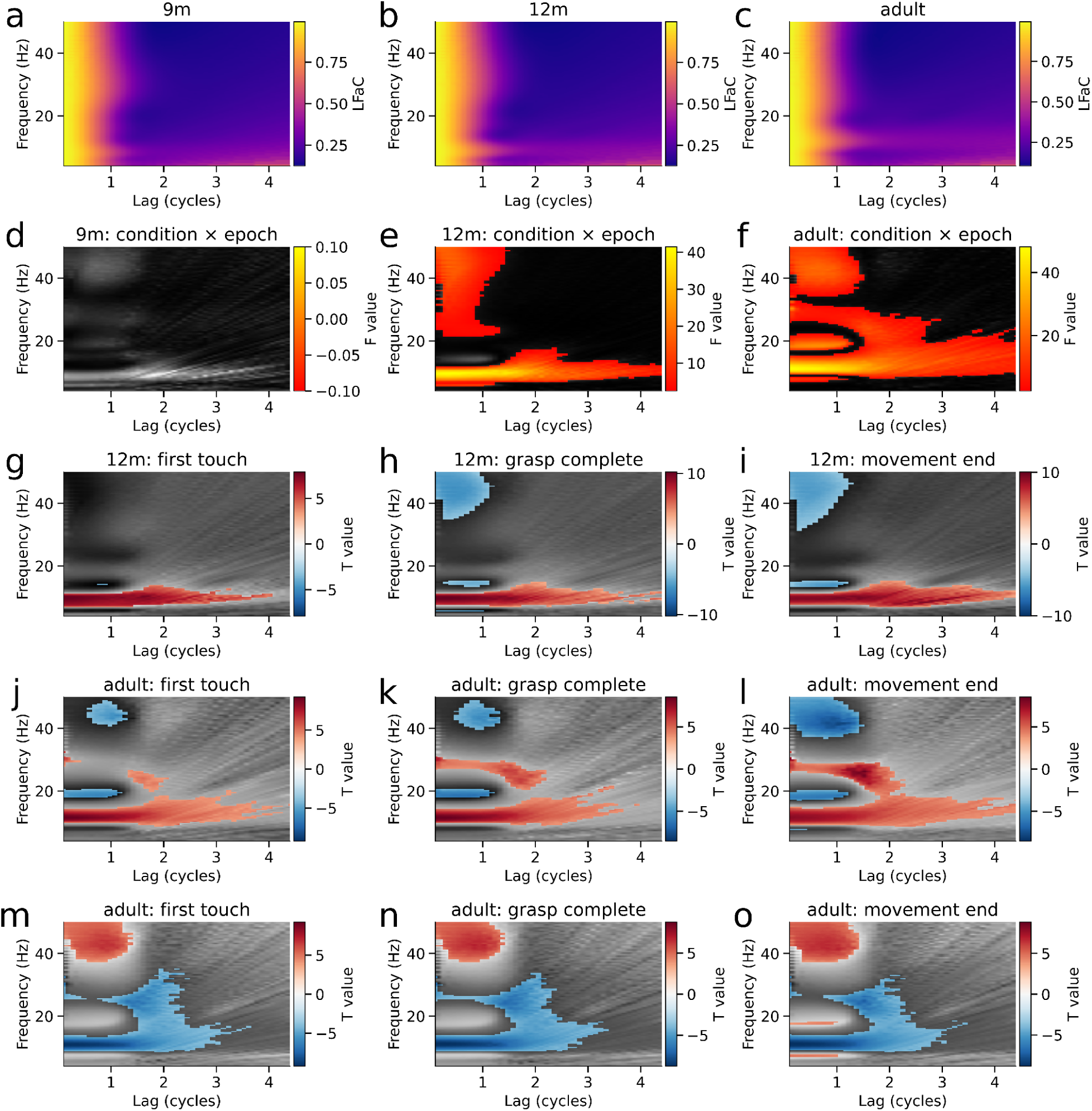
Lagged Fourier autocoherence (LFaC) reveals diffuse effects without frequency-specific structure. a-c) Grand-averaged LFaC within sensorimotor EEG clusters from 9-month infants (a), 12-month infants (b), and adults (c). d-f) F-statistics for the condition × epoch interaction from a cluster-based permutation ANOVA on LFaC values across participants in each group. Significant clusters (*p* < 0.05, corrected) are overlaid in color. g-i) T-statistics from cluster-based permutation tests comparing observation vs. execution for three movement-aligned epochs in 12-month infants. j-l) Corresponding tests for the adult group, showing scattered and broadband differences without spectral selectivity. m-o) Within-condition comparisons for the adult group, comparing each epoch to baseline, again showing diffuse activity without consistent spectral localization. In contrast to results obtained with LHaC (Figure 6), effects observed with LFaC do not exhibit clear frequency-band specificity.

To determine whether spectral rhythmicity differed between the observation and execution conditions across baseline and task-related epochs, a mass-univariate two-way repeated measures ANOVA was applied to LHaC in each group, with condition, epoch, and their interaction as factors (see Methods). In the 9-month-old group, no significant clusters were detected (Figure 6g), indicating no reliable modulation of rhythmicity by condition or epoch. In contrast, the 12-month-old group (Figure 6h) showed a significant condition-by-epoch interaction cluster in the gamma range, spanning 15.5-50.0 Hz and 0.1-1.0 lag cycles (*F*(1) = 31.96, *p* = 0.047). The adult group (Figure 6i) exhibited three significant clusters for this interaction: one in the theta range (4.0-7.5 Hz, 0.1-1.6 cycles; *F*(1) = 34.67, *p* = 0.034), a second in the alpha band (9.0-12.5 Hz, 0.1-4.4 cycles; *F*(1) = 41.33, *p* = 0.006), and a third in the gamma band (26.0-50.0 Hz, 0.1-1.6 cycles; *F*(1) = 21.55, *p* = 0.003). These results demonstrate that LHaC becomes increasingly sensitive to task-related rhythmic modulations with age, particularly in higher frequency bands. For comparison, the same ANOVA applied to LFaC detected no significant clusters in the 9-month group (Figure 7d), and only diffuse and overlapping clusters in the 12-month (5.0-18.0 Hz, 0.1-4.4 cycles and 21.0-50.0 Hz, 0.1-1.5 cycles; Figure 7e) and adult groups (6.5-32.5 Hz, 0.1-4.4 cycles; 16.5-22.0 Hz, 0.1-1.3 cycles; and 33.5-50.0 Hz, 0.1-1.5 cycles; Figure 7f), with less distinct frequency-band specificity.

We then followed up the significant condition × epoch interactions with pairwise comparisons between execution and observation conditions within each epoch. In the 12-month group, significant differences were observed across alpha, beta, and gamma bands (Figure 6j-l). During the first-touch epoch, alpha LHaC (∼6.5-8.0 Hz, 0.3-4.4 cycles) was lower during execution than observation (*t*(23) = 8.57, *p* = 0.001), while multiple clusters spanning 17-50 Hz and 0.1-0.8 cycles showed greater beta/gamma rhythmicity during execution (*t*(23) = 6.82-6.70, all *p* ≤ 0.008; Figure 6j). During the grasp-complete epoch, alpha again showed lower LHaC during execution (∼7-8 Hz, 0.2-4.1 cycles; *t*(23) = 8.63, *p* = 0.001), while a large cluster spanning 17-50 Hz and 0.1-0.8 cycles indicated increased beta/gamma coherence in execution (*t*(23) = 10.25, *p* = 0.001; Figure 6k). This pattern persisted into the movement-end epoch, with reduced alpha LHaC (∼7-8 Hz, 0.3-3.8 cycles; *t*(23) = 7.56, *p* = 0.001) and a large beta/gamma cluster (16.5-50 Hz, 0.1-0.8 cycles; *t*(23) = 8.62, *p* = 0.001) elevated in execution (Figure 6l). In adults (Figure 6m-o), similar condition differences were observed in the alpha band during all epochs (e.g., grasp-complete: 10.5-11.5 Hz, 0.2-1.7 cycles; *t*(21) = 6.39, *p* = 0.001; movement-end: 9.5-11.5 Hz, 0.1-2.4 cycles; *t*(21) = 8.75, *p* = 0.001), while theta LHaC (∼4-7 Hz, 0.1-1.7 cycles) was also greater in execution during the movement-end epoch (*t*(21) = 8.60, *p* = 0.001). In summary, whereas both 12-month-olds and adults exhibited reduced alpha rhythmicity during action execution relative to observation, 12-month-olds additionally showed increased beta and gamma coherence during execution, while adults uniquely showed enhanced theta rhythmicity following movement completion. In contrast, LFaC yielded a diffuse pattern of clusters that were less confined to specific frequency or lag bands (Figure 7g-l). Effects were broad and overlapping across theta, alpha, beta, and gamma ranges, with multiple clusters per epoch and substantial spread across lags.

We also performed follow-up pairwise comparisons between epochs within each condition. Notably, significant between-epoch differences were only observed in the execution condition of the adult group (Figure 6p-r). Across all three task-related epochs, alpha LHaC (∼9.5-11.5 Hz) was significantly reduced compared to baseline, with effects spanning lags from 0.2-2.6 cycles (first-touch: *t*(21) = 9.40, *p* = 0.001; grasp-complete: *t*(21) = 9.61, *p* = 0.001; movement-end: *t*(21) = 9.61, *p* = 0.001). In the same epochs, increases in gamma LHaC (33.5-50.0 Hz, 0.1-0.8 cycles) were observed (first-touch: *t*(21) = 7.39, *p* = 0.001; grasp-complete: *t*(21) = 7.93, *p* = 0.001; movement-end: *t*(21) = 7.93, *p* = 0.001), alongside more modest increases in beta (∼21 Hz) and theta (4.0-6.5 Hz) rhythmicity. These results indicate that during execution in adults, alpha coherence decreased while theta, beta, and gamma coherence increased in a task-phase-dependent manner. By contrast, LFaC revealed large and overlapping clusters within the same epochs (Figure 7m-o), with effects spanning most of the 8-35 Hz range and lags up to 3.8 cycles (e.g., movement-end: *t*(21) = 8.83, *p* = 0.001; 8.0-33.5 Hz, 0.1-3.8 cycles). These findings suggest that, in this dataset, LHaC yields more clearly delineated frequency-band-specific effects than LFaC when evaluating condition-dependent changes in rhythmicity.

Our simulations indicated that lower LHaC values reflect shorter oscillatory bursts in the frequency band of interest. To test whether the execution-related alpha-band LHaC reductions observed in infants and adults could reflect such changes in burst structure, we quantified alpha burst duration using a thresholded amplitude envelope approach. Linear mixed models revealed robust condition × epoch interactions for burst duration expressed both in cycles (9m: *χ*²(4) = 42.53, *p* < 0.001; 12m: *χ*²(4) = 693.84, *p* < 0.001; adults: *χ*²(4) = 987.59, *p* < 0.001) and in seconds (9m: *χ*²(4) = 75.47, *p* < 0.001; 12m: *χ*²(4) = 702.97, *p* < 0.001; adults: *χ*²(4) = 1037.11, *p* < 0.001). In cycles, 12-month-olds showed markedly shorter bursts during execution versus observation in the first-touch (*z* = -8.43, *p* < 0.001), grasp-completion (*z* = -11.84, *p* < 0.001), and movement-end epochs (*z* = -14.45, *p* < 0.001), with similar reductions in adults for grasp-completion (*z* = -9.17, *p* < 0.001) and movement-end (*z* = -13.99, *p* < 0.001) epochs. When expressed in milliseconds, the same pattern emerged: 12-month-olds exhibited shorter bursts in the first-touch (*z* = -7.71, *p* < 0.001), grasp-completion (*z* = -11.21, *p* < 0.001), and movement-end epochs (*z* = -13.53, *p* < 0.001), while adults again showed reductions in the same epochs (first-touch: *z* = -7.93; grasp-completion: *z* = -9.01; movement-end: *z* = -13.95; all *p* < 0.001). At 9 months, effects were more limited, with no significant execution–observation differences in these epochs. The convergence of the cycle-based and absolute duration analyses therefore supports the interpretation that execution-related decreases in alpha LHaC reflect shorter alpha bursts.

### 3.3 Alpha and beta rhythmicity index implicit sensorimotor adaptation over blocks of trials

To assess whether LHaC can detect dynamic, block-wise changes in neural rhythmicity during learning, we applied it to MEG data from a visuomotor adaptation task involving both implicit and explicit learning strategies (Szul et al., 2023). This dataset enabled us to evaluate three performance criteria of lagged autocoherence: (1) its alignment with spectral features (via RMSE to the PSD), (2) its sensitivity to gradual within-subject changes in oscillatory dynamics over learning, and (3) its ability to differentiate learning mechanisms across groups. We focused on alpha and beta bands given their established roles in sensorimotor control and adaptation, and used both visual and motor epochs to compare condition-dependent rhythmicity changes across task phases. By tracking crossover point and decay rate of inverse sigmoid fits to LHaC and LFaC, we quantified how rhythmicity evolved over time and between groups. This provided a direct test of whether LHaC could detect frequency-specific markers of implicit and explicit learning, serving as a test case for evaluating the temporal sensitivity and behavioral relevance of LHaC under realistic experimental noise and task complexity.

We computed LHaC and LFaC on MEG data from a cluster of sensors centered over the left sensorimotor cortex. Two groups of subjects performed joystick-based reaches with their right hand to visually presented targets after observing a random dot kinematogram with varying levels of coherent rotational motion in either the clockwise or counter-clockwise direction. In some trials, the position of the cursor controlled by the joystick was rotated 30 degrees relative to the actual movement of the joystick. For the implicit group, the rotation was always 30 degrees counter-clockwise and was unrelated to the direction of coherent motion in the RDK. These subjects could therefore compensate for the rotation using implicit sensorimotor adaptation (Taylor et al., 2014). For the explicit group, the direction of the rotation was entirely predicted by the direction of RDK coherent motion, and when there was no coherent motion there was no rotation. The optimal strategy for this group was therefore to use an explicit re-aiming strategy to compensate for the predicted perturbation (Taylor et al., 2014). After an initial training, each group completed one baseline block of trials, six blocks with visuomotor rotation, and one wash-out block.

Averaged across rotation blocks, the implicit and explicit groups exhibited comparable patterns of LHaC across frequencies and lags in both the visual (Figure 8a,b) and motor (Figure 8i,j) epochs. To evaluate how closely each lagged autocoherence metric aligned with spectral peaks, we computed normalized RMSE between the metric, averaged over 0.5-1 cycle, and the periodic component of the PSD: in the visual epoch, RMSE was 0.033 (LHaC) versus. 0.080 (LFaC) for the implicit group (Figure 8c) and 0.034 versus 0.079 for the explicit group (Figure 8d); in the motor epoch, RMSE was 0.042 (LHaC) versus 0.100 (LFaC) for the implicit group (Figure 8k) and 0.029 versus 0.093 for the explicit group (Figure 8l). To characterize dynamic changes in rhythmic activity across learning, we fit inverse sigmoid functions to LHaC and LFaC in the alpha and beta bands and examined changes in crossover point and decay rate across blocks, relative to the baseline block. In the visual epoch, significant group differences were observed in both the alpha and beta bands. The LHaC alpha crossover point differed in blocks 2-8 (all *p* < 0.032; Figure 8e), and decay rate in blocks 2-6 (all *p* < 0.026; Figure 8f). The LHaC beta crossover point differed in blocks 2-5 and 7 (all *p* < 0.038; Figure 8g), and decay rate in blocks 3 and 7 (all *p* < 0.042; Figure 8h). In the motor epoch, group differences in LHaC alpha crossover point and decay rate emerged in blocks 2-5 and 7 (all *p* < 0.045; Figure 8m–n), and beta showed similar differences in crossover point (blocks 2-4, 7; all p < 0.031) and decay rate (blocks 3, 7; all p < 0.021; Figure 8o-p). In each case, the implicit group showed earlier and more pronounced modulation. Except for alpha crossover in the visual epoch, all differences resolved in the washout block. In contrast, lagged coherence computed with LFaC revealed weaker and less consistent effects (Figure 9), suggesting that LHaC may be more effective in capturing frequency-specific learning-related changes.

**Figure 8.**
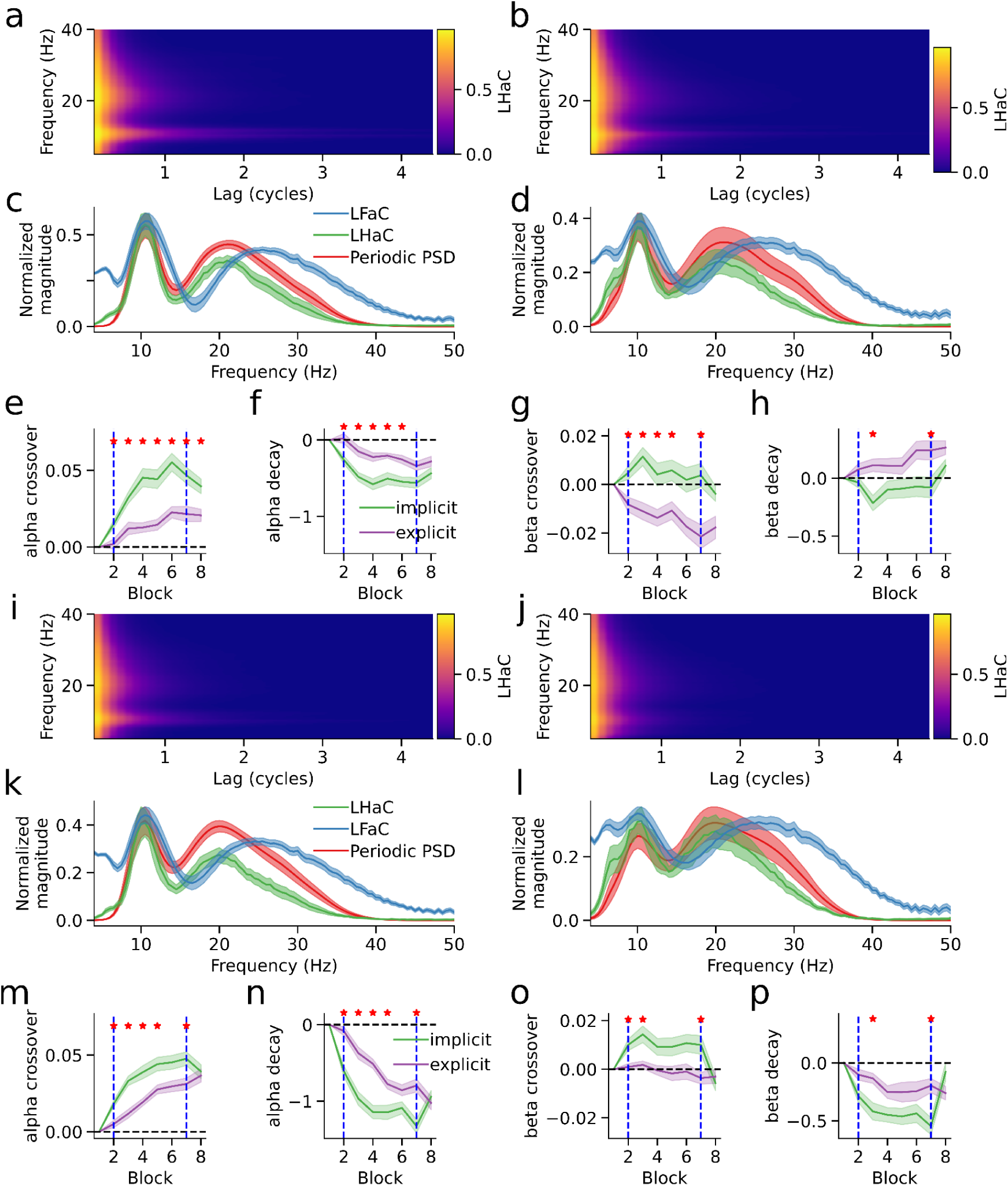
Lagged Hilbert autocoherence reveals dynamic modulation of rhythmic activity during visuomotor learning. a-b) Lagged Hilbert autocoherence in left central sensors for the visual epoch in the implicit (a) and explicit (b) groups, averaged over visuomotor rotation blocks. c-d) Normalized periodic PSD, LHaC, and LFaC (averaged over lags) for the visual epoch in the implicit (c) and explicit (d) groups. e-f) Alpha-band LHaC inverse sigmoid crossover point (e) and decay rate (f) for the visual epoch, shown across blocks. g-h) Beta-band LHaC crossover point (g) and decay rate (i) for the visual epoch. i-j) LHaC in left central sensors for the motor epoch in the implicit (i) and explicit (j) groups, averaged over visuomotor rotation blocks. k-l) Normalized periodic PSD, LHaC, and LFaC (averaged over lags) for the motor epoch in the implicit (k) and explicit (l) groups. m-n) Alpha-band crossover point (m) and decay rate (n), and o-p) beta-band crossover point (o) and decay rate (p) for the motor epoch. In panels e-h and m-p, solid lines show group means and shaded areas represent standard errors. Vertical dashed lines mark the start and end of visuomotor rotation blocks. Red stars indicate blocks where group differences were significant (p < 0.05, bootstrap permutation test).

**Figure 9.**
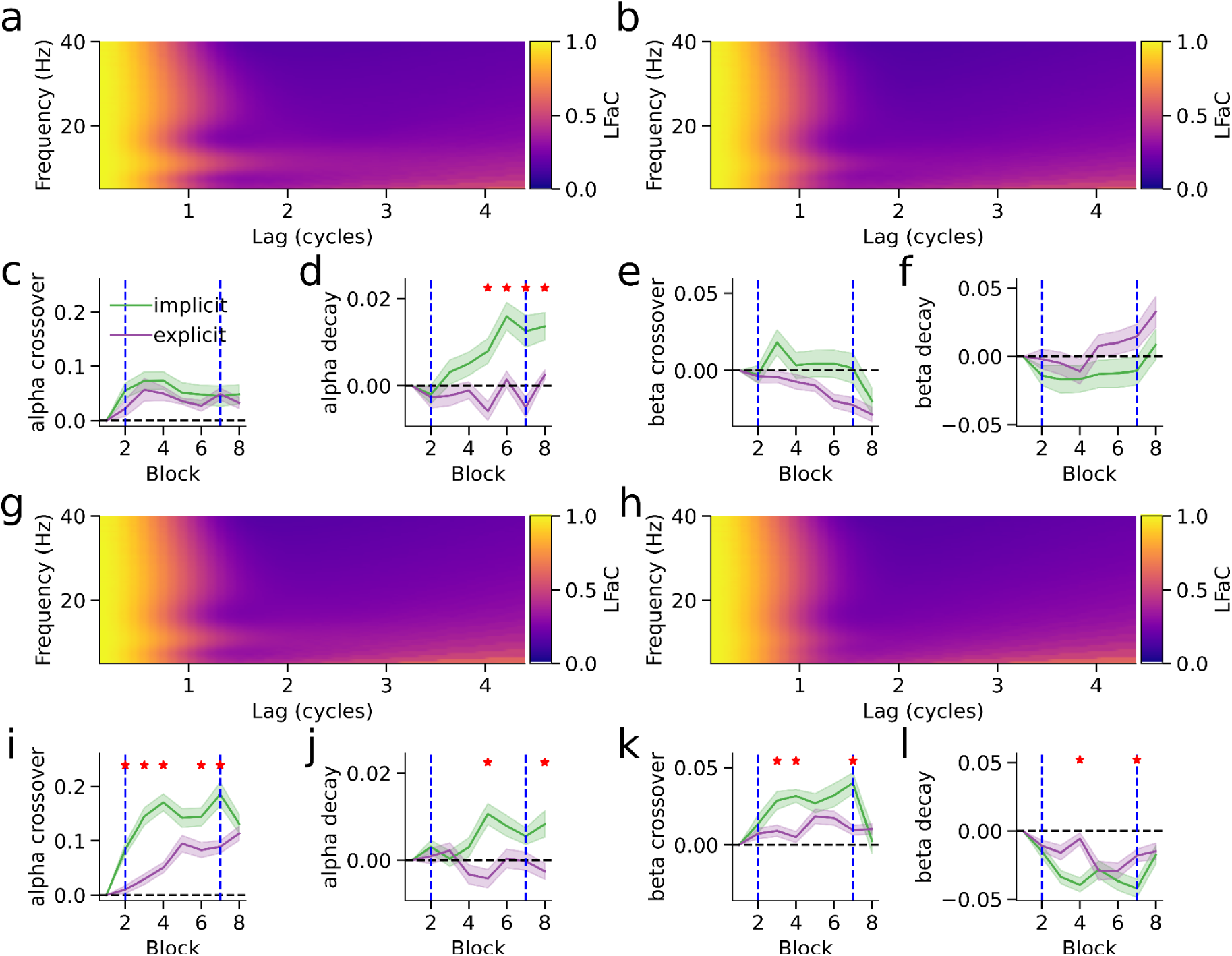
Lagged Fourier autocoherence is insensitive to dynamic modulation of rhythmic activity during visuomotor learning. a-b) Lagged Fourier autocoherence (LFaC) in left central sensors for the visual epoch in the implicit (a) and explicit (b) groups, averaged over visuomotor rotation blocks. c-d) Alpha-band LFaC inverse sigmoid crossover point (c) and decay rate (d) for the visual epoch, shown across blocks. e-f) Beta-band LFaC crossover point (e) and decay rate (f) for the visual epoch. g-h) LFaC in left central sensors for the motor epoch in the implicit (g) and explicit (h) groups. i-j) Alpha-band crossover point (i) and decay rate (j), and k-l) beta-band crossover point (k) and decay rate (l) for the motor epoch. In panels c-f and i-l, solid lines show group means and shaded areas represent standard errors. Vertical dashed lines mark the start and end of visuomotor rotation blocks. Red stars indicate blocks where group differences were significant (p < 0.05, bootstrap permutation test).

We followed up the bootstrapped permutation tests with linear mixed models, which allowed us to assess not only group and block-type differences but also linear trends across the six visuomotor rotation blocks. In the visual epoch, LHaC revealed robust and condition-specific modulation of rhythmic activity during sensorimotor learning. For the crossover point, there was a significant group × block-type interaction in the alpha band (*χ*²(2) = 6.87, *p* = 0.032), with the implicit group exhibiting significant increases from baseline to both rotation and washout (both *p* < .0001), whereas the explicit group showed no change between blocks; both groups, however, showed a linear increase across the rotation blocks (*χ*²(1) = 27.60, *p* < 0.001). Alpha decay rate had a main effect of block-type (*χ*²(2) = 22.32, *p* < 0.001), decreasing from baseline to rotation (*t*(220) = 5.51, *p* < 0.001), within the rotation blocks (*χ*²(1) = 12.85, *p* < 0.001), and from rotation to washout (*t*(220) = 4.70, *p* < 0.001). Beta crossover point also showed a group × block-type interaction (*χ*²(2) = 6.38, *p* = 0.041), with an increase from baseline to rotation only in the implicit group (*t*(220) = 2.41, *p* = 0.044), and a linear decrease within rotation blocks (*χ*²(2) = 4.87, *p* = 0.027). Beta decay rate had a main effect of block type (*χ*²(2) = 7.76, *p* = 0.021), though follow-up comparisons were not significant. In contrast, LFaC in the visual epoch showed only main effects of block type or group in alpha and beta, with some additional group × block-type interactions and main effects of block within rotation.

In the motor epoch, LHaC again revealed pronounced condition-specific modulations. For the alpha crossover point, there was a significant group × block-type interaction (*χ*²(2) = 73.50, *p* < 0.001): both groups increased from baseline to rotation (both *p* < 0.001), linearly increased during the rotation blocks (χ²(1) = 67.35, p < 0.001), and the explicit group showed a further increased from rotation to washout (*t*(220) = 2.84, *p* = 0.014). Alpha decay rate showed a group × block-type interaction (*χ*²(2) = 17.48, *p* < 0.001), decreasing from baseline to rotation (both *p* < 0.001), and linearly during rotation (*χ*²(1) = 83.59, *p* < 0.001) in both groups, and further decreasing from rotation to washout in the explicit group (t(220) = 3.69, p < 0.001). For the beta crossover point, there was a group × block-type interaction (*χ*²(2) = 9.24, *p* = 0.010), with the implicit group increasing from baseline to rotation (*t*(220) = 2.90, *p* = 0.012), and decreasing from rotation to washout (*t*(220) = 4.46, *p* < 0.001), returning to baseline levels. Beta decay rate also showed a group × block-type interaction (*χ*²(2) = 11.36, *p* = 0.003), decreasing from baseline to rotation (*t*(220) = 4.84, *p* < 0.001), and increasing from rotation to washout (*t*(220) = 3.97, *p* < 0.001) in the implicit group only, with a linear decrease during the rotation blocks in both groups (*χ*²(1) = 5.36, *p* = 0.021). LFaC in the motor epoch again showed generally weaker effects: alpha exhibited a group × block-type interaction along with main effects of group and block within rotation, while beta showed a group × block-type interaction and main effects of block type and block within rotation.

Overall, across both visual and motor epochs, LHaC consistently revealed stronger and more specific group- and block-related modulations in both alpha and beta rhythms compared with LFaC, particularly in detecting interaction effects and linear trends across the visuomotor rotation blocks. Taken together, these results indicate that LHaC captures clear and behaviourally relevant changes in oscillatory dynamics during sensorimotor adaptation, with alpha and beta rhythms showing distinct temporal profiles across learning and washout. Alpha activity tended to shift its crossover point upward and its decay rate downward over the course of rotation. Beta changes were more condition-specific, with the implicit group showing stronger modulation between baseline and rotation, and return to baseline levels during washout, especially in the motor epoch. Compared with LFaC, LHaC was more sensitive to interaction effects and to gradual changes across the visuomotor rotation blocks, suggesting it may be a better tool for tracking fine-grained learning-related dynamics.

## 4. Discussion

In this paper we introduced a novel algorithm, lagged Hilbert autocoherence (LHaC), to characterize the rhythmicity of neuronal oscillations. As a refinement of lagged Fourier autocoherence (LFaC; Fransen et al., 2016, 2015), LHaC retains the conceptual foundation of measuring coherence at increasing temporal lags while addressing key limitations of LFaC, including poor spectral resolution, aliasing at higher frequencies, and conflation with amplitude covariation. By using precise frequency-domain filtering and Hilbert-based analytic signals, LHaC produces lagged autospectra with sharper, more spectrally accurate peaks that better match the periodic component of power spectral densities (PSDs). Simulation results demonstrated that LHaC exhibited closer empirical correspondence to the spectral profile of the simulated signal and greater spectral resolution under finite, noisy conditions. These differences reflect distinct mathematical formulations, rather than superiority in estimating a shared population quantity. More generally, any time-frequency representation, whether Fourier, Hilbert, wavelet, or otherwise, can be used to define its own lagged autocoherence function, each with different theoretical underpinnings. Our comparisons are therefore descriptive of the empirical behavior of the estimators under simulation and experimental conditions, not formal evaluations of statistical performance. We further showed that these empirical advantages make LHaC useful for revealing condition- and learning-related differences in neural rhythmicity, particularly in alpha and beta bands during action observation and execution.

Using EEG data, we found developmental increases in both alpha peak frequency and rhythmicity from infancy to adulthood, replicating prior findings using LFaC (Rayson et al., 2023, 2022) but with improved spectral resolution. In all age groups, alpha rhythmicity was lower during action execution than observation, consistent with prior work on alpha power modulations during motor tasks (Aleksandrov and Tugin, 2012; Debnath et al., 2019). Consistent with our simulation results, direct analysis of alpha burst structure confirmed that these execution-related LHaC reductions were accompanied by significantly shorter and more transient alpha bursts in 12-month-olds and adults, but not in 9-month-olds (Schaworonkow and Voytek, 2021). Critically, LHaC revealed frequency-specific differences in task-related rhythmicity that were less distinct in LFaC: in 12-month-olds and adults, execution was associated with increased beta and gamma rhythmicity, particularly toward the end of the movement, consistent with the developmental emergence of the post-movement beta rebound (Cassim et al., 2000; Cheyne, 2013; Jurkiewicz et al., 2006) and movement-related gamma synchrony (Cheyne et al., 2008; Muthukumaraswamy, 2010; Gaetz et al., 2020, 2010; Hao et al., 2019). By contrast, epoch-related changes in LHaC were absent in 9-month-olds, suggesting reduced task modulation at this age. Importantly, the frequency-band specificity and temporal structure of these effects were not captured by LFaC, which yielded broader and less differentiated clusters. These findings highlight LHaC’s utility in detecting developmentally emergent and condition-specific changes in neural rhythmicity, and suggest that it may be more sensitive than power or Fourier-based coherence to age-dependent changes in oscillatory structure.

In the adult MEG dataset involving a sensorimotor adaptation task, LHaC revealed robust, frequency-specific modulation of rhythmicity in both visual and motor epochs that was less distinct in LFaC. In the visual epoch, alpha rhythmicity increased across rotation blocks in both groups, with a steeper rise in the implicit group, while beta rhythmicity increased from baseline to rotation only in the implicit group before declining during washout. In the motor epoch, both groups showed increased alpha rhythmicity during rotation, but the implicit group exhibited an earlier and sharper rise, whereas beta rhythmicity again increased selectively in the implicit group during rotation and returned to baseline in washout. Across both epochs, these learning-related changes were more temporally specific and statistically robust in LHaC than in LFaC, which yielded weaker and less consistent patterns. Linking these empirical findings to our simulations, the observed increases in crossover point and decreases in decay rate in alpha and beta bands during adaptation are consistent with longer burst durations, suggesting that visuomotor learning enhances the temporal stability of oscillatory events in sensorimotor networks. The early modulation of alpha rhythmicity may reflect more rapid engagement of general error-processing mechanisms (Driel et al., 2012; Navarro-Cebrian et al., 2013), whereas the observed changes in beta rhythmicity align with prior findings that implicate beta activity as a marker of implicit sensorimotor adaptation (Jahani et al., 2020), possibly mediated by bursts with distinct waveform shapes during the beta rebound period (Szul et al., 2023). Across both epochs, these learning-related changes were more temporally specific and statistically robust in LHaC than in LFaC, which yielded weaker and less consistent patterns.

Whilst lagged autocoherence discards the assumption of phase consistency over time, it still assumes stationarity in peak frequency. LHaC and related methods, therefore, cannot detect rhythmic signals that change in peak frequency. However, we know that there are changes in instantaneous frequency in some signals (Benwell et al., 2019; Nelli et al., 2017) which are difficult to track using methods based on bandpass filtering. Variants of empirical mode decomposition have been proposed to separate neural signals into modes which may vary in instantaneous frequency (Fabus et al., 2021; Huang et al., 1998; Quinn et al., 2021b, 2021a), thereby isolating signal sources that are mixed together in sensor signals. An interesting future direction would therefore be to adapt LHaC to apply to each mode derived from EMD in order to examine changes in rhythmicity of frequency-varying neural activity. Considering significance testing, we note that our current use of phase-randomized surrogates to threshold low-SNR estimates implicitly assumes that the original signal is both stationary and Gaussian. While this assumption is reasonable in many contexts, it may be violated in neural time series that exhibit strong nonstationarity or heavy-tailed amplitude distributions. To address this limitation, future work could explore alternative surrogate generation methods, such as time-frequency permutation or model-based residual bootstrapping, that preserve key statistical properties of the data while eliminating lagged structure. Such approaches may yield more accurate and robust thresholds for detecting meaningful coherence in neural signals. Finally, it is important to emphasize that our analyses compare the behavior of two distinct estimators rather than assessing their accuracy with respect to a shared population-level lagged autocoherence function. Because we did not specify a generative time-series model with a defined population lagged autocoherence function, the simulated data lack a formal ground truth against which estimator bias or variance can be assessed. Future work could complement our empirical approach with model-based simulations, allowing for a parametric definition of lagged autocoherence and a more classical evaluation of estimator properties.

One of the strengths of our approach is spectral precision, which results from narrow bandpass filtering. However, this also reflects a potential weakness: in frequency ranges with low power, narrow bandpass filtering introduces amplitude correlations which result in spuriously high lagged autocoherence. Another approach could be to focus exclusively on the phase, using lagged phase-locking value (Bruña et al., 2018; Lachaux et al., 2000). This approach was used in a recent study (Myrov et al., 2024), whereby ‘rhythmicity’ was quantified using a phase-autocorrelation function (pACF) derived from the phase-locking value. However, this necessitated a correction of the signal lag using the mean instantaneous frequency of the narrow-band signal to avoid a bias towards higher frequencies. In contrast, we used a threshold computed from phase-shuffled surrogate data, which has the advantage that the metric is thus adapted to the SNR of the dataset (Figure S2).

The recent surge in sophisticated techniques for analyzing frequency-specific neural activity has been largely fueled by a growing appreciation of the potential for activity in some frequency ranges, most notably beta and gamma, to manifest not as continuous oscillations, but as discrete, transient bursts (Feingold et al., 2015; Lundqvist et al., 2016; Sherman et al., 2016). The validity of this notion could radically reshape theories anchored in the concept of phase precession of oscillatory activity being perpetuated over extended time intervals. Conversely, the appearance of burst-like activity could be an artifact of our recording methods, only made visible when the amplitude crosses a certain detectability threshold, with the actual activity being driven by an underlying rhythmic process (van Ede et al., 2018). Lagged autocoherence has previously been used to discern whether activity within a specific frequency band is “bursty” or oscillatory (Fransen et al., 2016; Little et al., 2019; Rayson et al., 2023, 2022; Szul et al., 2023), but detection of bursts has predominantly relied on power measurements (Brady and Bardouille, 2022; Rayson et al., 2022; Shin et al., 2017; Szul et al., 2023; though see Cole and Voytek, 2019). Future work could explore a temporally-resolved version of lagged Hilbert autocoherence to detect frequency-specific, oscillatory bursts with greater precision. This might involve computing LHaC at each time-point with a short, fixed-cycle lag (e.g., 0.5 cycles), potentially in combination with wavelet-based filtering to optimize the time-frequency tradeoff. Such an approach could enable fine-grained tracking of burst onset, offset, and rhythmicity, offering a complementary alternative to threshold-based burst detection.

Algorithms that effectively parameterize the periodic and aperiodic components of power spectral densities (Barry and Blasio, 2021; Donoghue et al., 2020; Gerster et al., 2022; Wen and Liu, 2016) have sparked significant advances in systems and cognitive neuroscience, transitioning the field away from the conventional tendency to conflate these two distinct signal sources. Benefiting from these algorithms, numerous studies have found that task-specific modulation of aperiodic activity can provide crucial insight into cognitive processing (Waschke et al., 2021), shifts in the slope of aperiodic activity can index excitation/inhibition balance (Gao et al., 2017), and abnormal periodic and aperiodic activity can serve as early indicators of neurodegenerative diseases (Karalunas et al., 2022; Ostlund et al., 2021). Despite their evident value, these methodologies rely on the presumption that neural signals can be cleanly partitioned into rhythmic (periodic) and arhythmic (aperiodic) components based solely on the shape of the power spectral density (PSD). They thus operate under the assumption that a 1/f model of aperiodic activity can be robustly fit to the PSD, and then periodic activity can be parameterized from the residuals of this fit. Two key advantages of LHaC could be leveraged in future work to simultaneously fit both the aperiodic and periodic components of the PSDs. The first advantage is the method’s high spectral accuracy and resolution, which can match that of the PSD. The second advantage is that, at short lags, lagged Hilbert autocoherence is, by definition, proportional to the periodic component of the PSD. While speculative, we suggest that integrating LHaC into peak-fitting routines may help reduce the number of missed periodic peaks and spurious high frequency peaks, potentially leading to more accurate and robust spectral parameterization.

## 5. Conclusion

In conclusion, this study introduces lagged Hilbert autocoherence (LHaC) as a refined estimator for assessing neural rhythmicity, addressing limitations of Fourier-based approaches such as spectral imprecision, aliasing, and amplitude confounds. Through simulations and empirical applications to EEG and MEG datasets, we demonstrate that LHaC yields more spectrally resolved and empirically stable estimates of rhythmic structure, enabling clearer detection of condition-, age-, and learning-related changes in alpha and beta activity. While LHaC and LFaC estimate distinct population quantities, the empirical advantages of LHaC in noisy, finite signals make it a valuable addition to the analytical toolbox for studying neural oscillatory dynamics.

## Data and Code Availability

Implementations of both LFaC and LHaC in MATLAB and python, demos, and code for all simulations and analyses can be found at https://github.com/danclab/lagged_hilbert_autocoherence. The data that support the findings of this study are available from the corresponding author, H.R., upon reasonable request.

## Author Contributions

S.Z.: Conceptualization, Methodology, Software, Formal analysis, Writing - Original Draft, Visualization. M.J.S: Methodology, Software, Validation. S.P.: Methodology, Software, Validation. A.M.: Formal analysis, Software, Validation. H.R: Conceptualization, Methodology, Writing - Original Draft, Writing - Review & Editing, Supervision. J.J.B. - Conceptualization, Methodology, Software, Formal analysis, Writing - Original Draft, Writing - Review & Editing, Supervision, Visualization, Funding acquisition.

## Supplementary Figures

**Figure S1.**
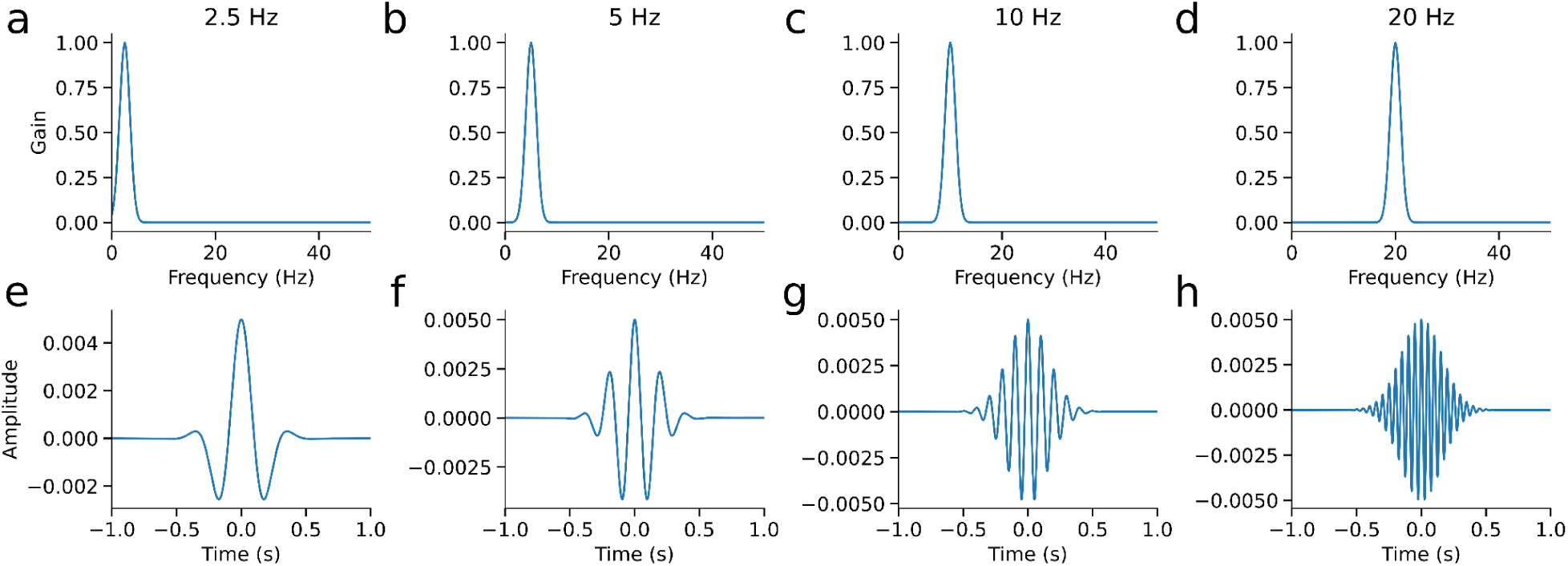
Frequency and impulse responses of Gaussian filters used for frequency-domain bandpass filtering. a-d) Frequency responses of Gaussian kernels centered at 2.5, 5, 10, and 20 Hz, respectively, each with fixed spectral resolution (σ = df/2, where df = 2 Hz). e-h) Corresponding time-domain impulse responses, obtained by inverse Fourier transform of the Gaussian frequency kernels.

**Figure S2.**
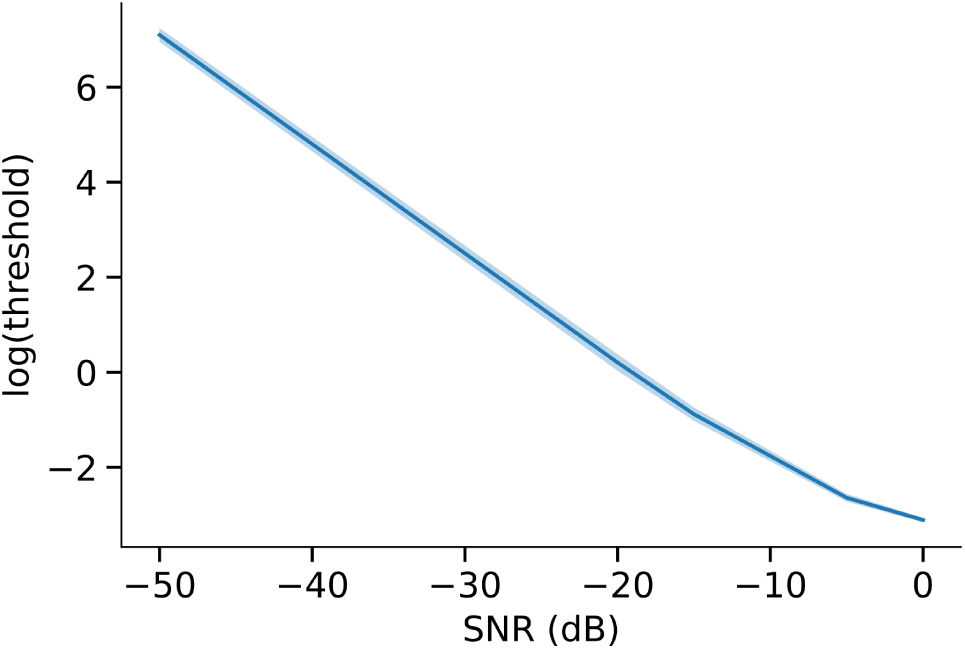
Relationship between signal-to-noise ratio (SNR) and the surrogate threshold for the joint amplitude normalization factor used in lagged Hilbert autocoherence (LHaC). Simulated signals consisted of 15 Hz bursts embedded in pink noise at varying SNR levels. For each SNR, the 95th percentile of the joint amplitude normalization factor was computed from 1000 phase-shuffled surrogates. Shown are the mean and standard deviation of the log-transformed thresholds across trials. As SNR increases, the surrogate threshold decreases, reflecting reduced amplitude normalization in signals with stronger oscillatory components.

**Figure S3.**
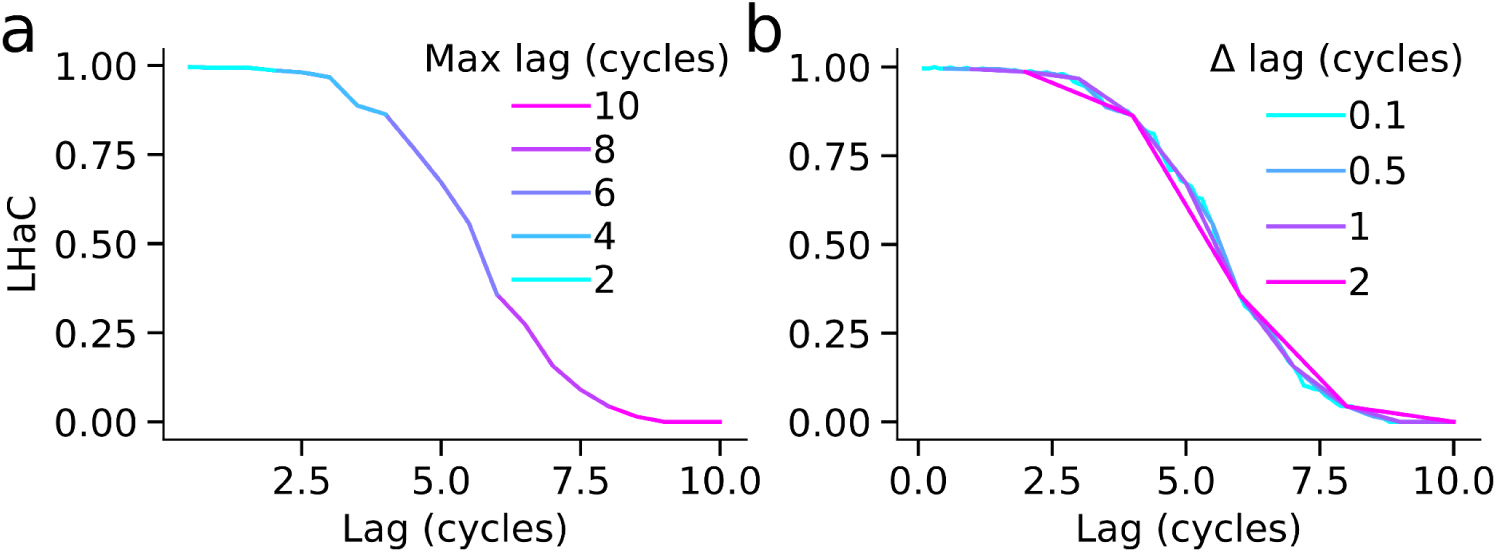
Sensitivity of lagged Hilbert autocoherence (LHaC) to lag sampling parameters. Simulated data contained transient 15 Hz bursts embedded in 1/f noise. a) Mean LHaC at 15 Hz as a function of lag duration for five maximum lag values (2-10 cycles), with lag increments fixed at 0.5 cycles. Increasing the maximum lag reveals the timescale over which coherence persists. b) Mean LHaC at 15 Hz for different lag sampling intervals (Δ lag = 0.1-2 cycles), with the maximum lag fixed at 10 cycles. While finer increments yield smoother LHaC curves, the overall coherence profile remains stable across settings, indicating that LHaC is robust to reasonable variation in lag sampling parameters.

**Figure S4.**
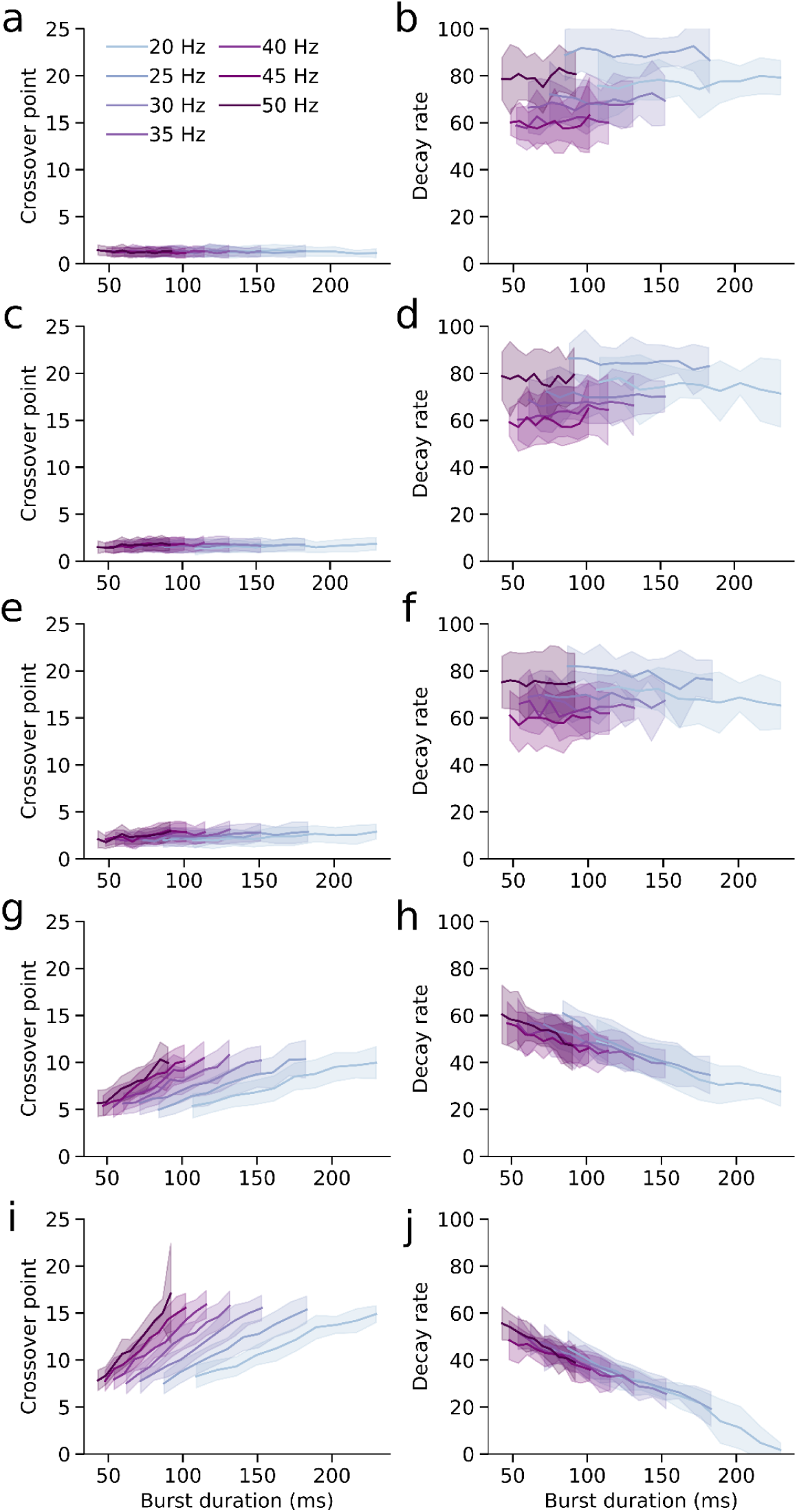
Lagged Hilbert autocoherence (LHaC) reflects burst duration across signal-to-noise ratios (SNRs). a-j) At each SNR level (-50, -20, -15, -5, and 0 dB, top to bottom), we evaluated the relationship between burst duration in milliseconds (x-axis) and the fitted parameters of an inverse sigmoid model applied to LHaC: the crossover point (left column) and decay rate (right column). Each line represents a different simulated frequency (20-50 Hz), with shaded areas indicating ±1 SD across trials. As SNR increases, both the crossover point and decay rate show progressively stronger and more monotonic relationships with burst duration across frequencies.

**Figure S5.**
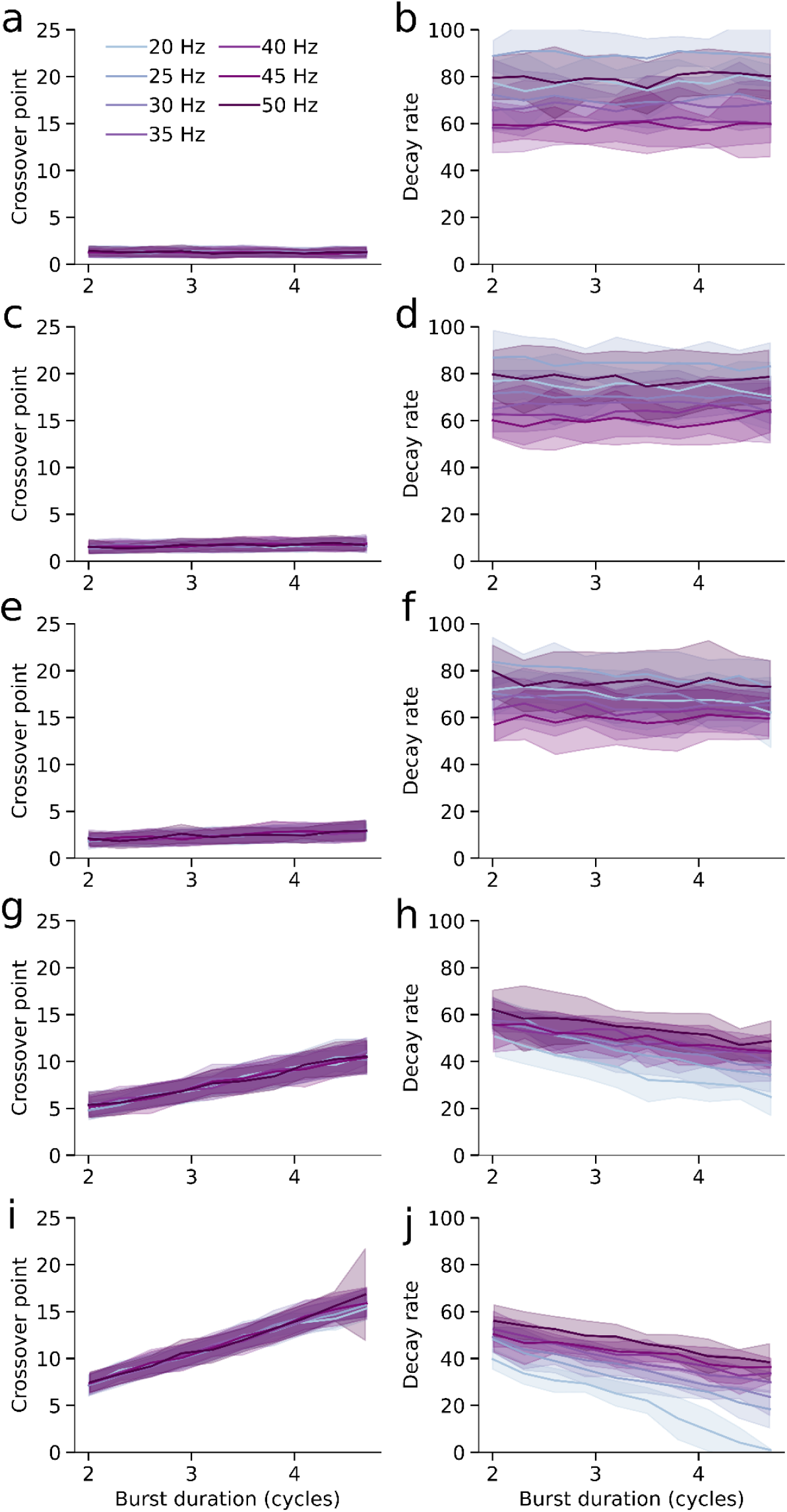
Lagged Hilbert autocoherence (LHaC) reflects burst duration in cycles across signal-to-noise ratios (SNRs). a-j) As in Figure S4, we examined the relationship between burst duration and LHaC parameters, but here duration is expressed in cycles rather than milliseconds. Each subplot pair shows the crossover point (left) and decay rate (right) of the inverse sigmoid fit to LHaC, plotted as a function of burst duration in cycles at an SNR of -50, -20, -15, -5, and 0 dB (top to bottom).

**Figure S6.**
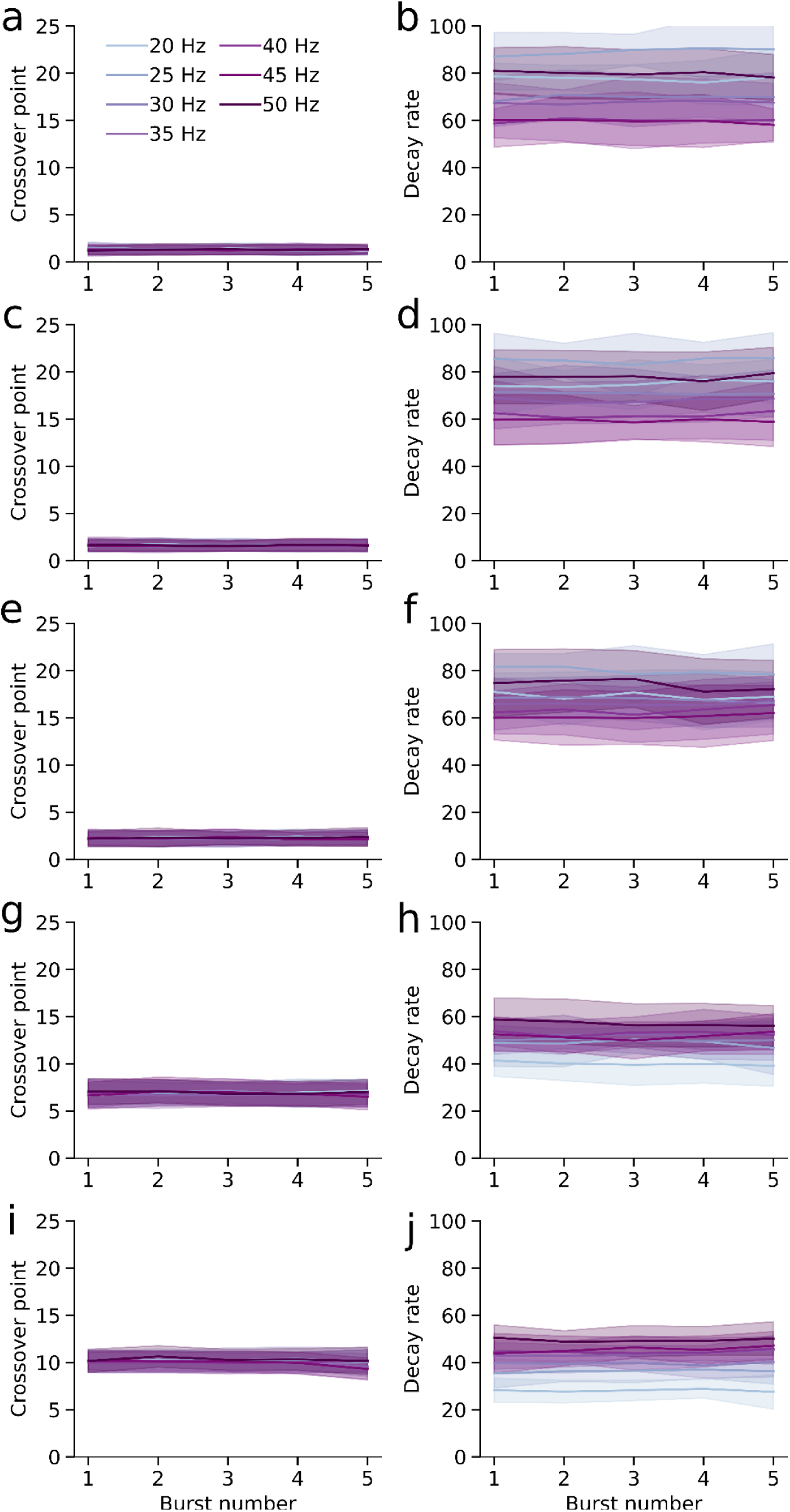
Lagged Hilbert autocoherence (LHaC) is invariant to burst count across signal-to-noise ratios (SNRs). a-j) As in Figure S4, we examined the relationship between burst number and LHaC parameters. Each subplot pair shows the crossover point (left) and decay rate (right) of the inverse sigmoid fit to LHaC, plotted as a function of burst number at an SNR of -50, -20, -15, -5, and 0 dB (top to bottom).

